# Regulation of angiogenesis by endocytic trafficking mediated by cytoplasmic dynein 1 light intermediate chain 1

**DOI:** 10.1101/2024.04.01.587559

**Authors:** Dymonn Johnson, Sarah Colijn, Jahmiera Richee, Joseph Yano, Margaret Burns, Andrew E. Davis, Van N. Pham, Amra Saric, Akansha Jain, Ying Yin, Daniel Castranova, Mariana Melani, Misato Fujita, Stephanie Grainger, Juan S. Bonifacino, Brant M. Weinstein, Amber N. Stratman

## Abstract

Dynein cytoplasmic 1 light intermediate chain 1 (LIC1, *DYNC1LI1*) is a core subunit of the dynein motor complex. The LIC1 subunit also interacts with various cargo adaptors to regulate Rab-mediated endosomal recycling and lysosomal degradation. Defects in this gene are predicted to alter dynein motor function, Rab binding capabilities, and cytoplasmic cargo trafficking. Here, we have identified a *dync1li1* zebrafish mutant, harboring a premature stop codon at the exon 12/13 splice acceptor site, that displays increased angiogenesis. *In vitro*, LIC1-deficient human endothelial cells display increases in cell surface levels of the pro-angiogenic receptor VEGFR2, SRC phosphorylation, and Rab11-mediated endosomal recycling. *In vivo*, endothelial-specific expression of constitutively active *Rab11a* leads to excessive angiogenesis, similar to the *dync1li1* mutants. Increased angiogenesis is also evident in zebrafish harboring mutations in *rilpl1/2*, the adaptor proteins that promote Rab docking to Lic1 to mediate lysosomal targeting. These findings suggest that LIC1 and the Rab-adaptor proteins RILPL1 and 2 restrict angiogenesis by promoting degradation of VEGFR2-containing recycling endosomes. Disruption of LIC1- and RILPL1/2-mediated lysosomal targeting increases Rab11-mediated recycling endosome activity, promoting excessive SRC signaling and angiogenesis.

## INTRODUCTION

Angiogenesis, the formation of new blood vessels from pre-existing vessels, is critical for sustained nutrient delivery during embryogenesis, homeostasis, and tumorigenesis. Angiogenesis is driven by cell signals such as those from the vascular endothelial growth factor (VEGF) family, as well as by physical changes in cell morphology and motility (1, 2). Endosomal trafficking plays both positive and negative roles in the regulation of pro-angiogenic signaling in endothelial cells, which are regulated in part by intracellular motor complexes (3–8).

The cytoplasmic dynein-dynactin protein complex is responsible for microtubule minus-end-directed movement of endosomes in eukaryotic cells (9–11). The 1.5 megadalton dynein complex is composed of six different subunits that are each dimerized in the full complex: two heavy chains (HCs) with the catalytic motor domain, two intermediate chains (ICs), six light chains (LCs), and two light intermediate chains (LICs) (9–11). Despite the critical nature of this motor complex, the role of LICs in particular has been understudied across all systems. LIC1, and LIC2 are encoded by two genes, *DYNC1LI1* and *DYNC1LI2,* respectively. LIC1 and LIC2 have been implicated in mitosis (12, 13), vesicular cargo binding (14), and lysosome and late endosome maturation (15). Aberrant expression of *DYNC1LI1* has also been linked with pancreatic ductal adenocarcinoma, colon cancer, hepatocellular carcinoma, and prostate cancer (16, 17), underscoring the need to better understand LIC function. However, the role of dynein motor components such as LIC1 in regulating the endothelium, and in particular developmental angiogenesis, remains entirely unknown.

In the endo-lysosomal pathway, cargoes are internalized from the plasma membrane inside early endosomes and retrogradely transported on microtubules via the dynein-dynactin motor complex. Internalized cargoes are subsequently either recycled back to the plasma membrane or delivered to late endosomes and lysosomes for degradation (18–23). Over the course of these movements, endosomes associate with different Rab GTPases. Rab4/Rab11a association promotes recycling back to the plasma membrane, while Rab7/Rab36 association promotes maturation to late endosomes and targeting to lysosomes (21, 22, 24–26). Adaptor proteins have been identified that facilitate interactions between Rab proteins and the dynein motor complex, including RILP (Rab Interacting Lysosomal Protein), RILPL1, and RILPL2 (Rab Interacting Lysosomal Protein Like 1 and 2). These interactions involve Rab docking to LIC1, promoting lysosomal targeting of signaling receptors (27–29) and consequent signal termination. VEGF signaling is a key contributor to angiogenesis (30), as well as vascular permeability, vascular homeostasis, endothelial cell (EC) proliferation, migration, arterial/venous identity, survival, and junctional stability (31). VEGF receptors transduce intracellular signals through a variety of mediators, including the phosphotidylinositol-3 kinase (PI3K)/AKT pathway, the Src family of non-receptor tyrosine kinases (SRC), and the extracellular signal-regulated kinases (ERK1/2) via the phospholipase C (PLC)/protein kinase C (PKC) pathway (32, 33). VEGF signaling is regulated in part by endosomal trafficking of the VEGF-VEGFR2 (vascular endothelial growth factor receptor 2) complex (3, 34). Upon VEGF binding, VEGFR2 dimerization induces receptor autophosphorylation and internalization to activate downstream signaling via ERK1/2 and AKT phosphorylation (35). Signal transduction strength and duration through the VEGF/VEGFR2 pathway can be modulated by altering the balance of endocytic recycling versus degradation (31, 35). Additionally, some VEGFR2-mediated signaling activation, i.e. via SRC, can occur at the plasma membrane (31, 36, 37). Therefore, the correct balance between endosome-to-plasma membrane recycling and lysosomal targeting of pro-angiogenic receptors is critical for proper vascular formation and homeostasis (20, 31, 38). The ability to promote vessel growth or regression has enormous therapeutic implications for several vascular-related pathologies including cancer, diabetes, stroke, and atherosclerosis (39).

In this study, we investigate the angiogenic phenotypes of a novel mutation identified in zebrafish *dynein cytoplasmic 1 light intermediate chain 1* (*dync1li1)* and how it alters endocytic trafficking, endothelial cell activation, migration, and VEGFR2 receptor availability. Our findings suggest that the C-terminal domain of LIC1 is critical for RILPL1/2 adaptor binding and for Rab-targeted lysosomal targeting. In the absence of the Lic1 C-terminal domain or RILPL1/2, endothelial cells (EC) over-recycle pro-angiogenic cargo back to the cell surface to promote excess angiogenesis.

## RESULTS

### LIC1 limits angiogenic sprouting and cellular motility *in vitro* and *in vivo*

We identified a novel zebrafish mutant allele (*dync1li1^y151^*) from a forward genetic ENU mutagenesis screen carried out on the *Tg(fli:eGFP)^y1/y1^*background that was designed to identify zebrafish mutants with vascular patterning defects (40). By 96 hours post-fertilization (hpf), *dync1li1^y151/y151^* mutant animals exhibit aberrant ectopic blood vessel sprouting from the intersegmental vessels (ISVs; **Fig. 1A,C,D**) and sub-intestinal vessels (SIVs; **Fig. 1A,E,F**) compared to their *dync1li1^WT^* siblings, with approximately 87% of ISVs displaying at least one ectopic sprout or branch (**Fig. 1B**). Fine mapping and positional cloning revealed that the causative mutation is a 19-base pair (bp) insertion at the exon 12/13 splice acceptor site in the *dync1li1* transcript, which introduces a premature stop codon predicted to result in a C-terminally truncated protein (**Supp. Fig. 1A**). In addition to their vascular defects, homozygous mutant animals have smaller eyes and a curved body axis. (**Fig. 1G**).

**Figure 1:**
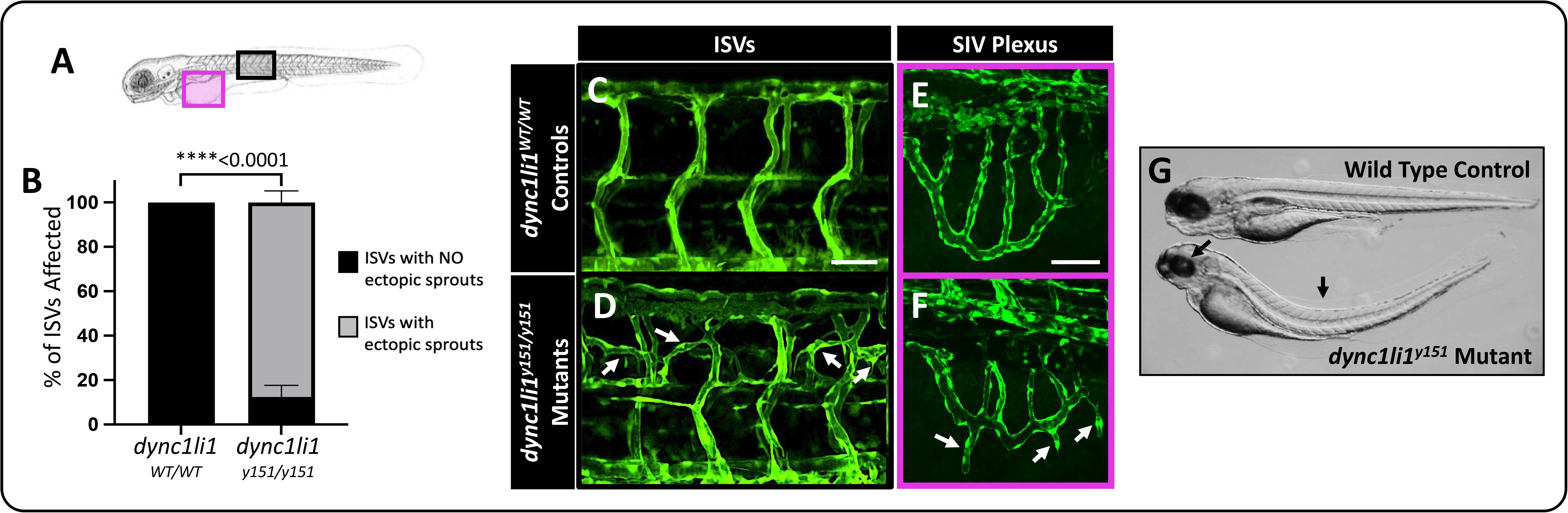
*dync1li1^y151^* mutant zebrafish exhibit excessive angiogenesis. **A.** Schematic demonstrating imaged areas in panels B-E. These areas include intersegmental vessels (ISVs, black box), and the sub-intestinal vascular plexus (SIVs, magenta box). **B.** Percentage of ISVs displaying ectopic sprouts in *dync1li1^WT/WT^* controls and *dync1li1^y151/y151^*mutants at 96 hpf. n=4 embryos, 10-15 ISVs analyzed per embryo. **C-F.** Confocal images of ISVs (C,D) or SIVs (E,F) from *dync1li1^WT/WT^* control (C,E) and *dync1li1^y151/y151^* mutant siblings (D,F) at 96 hpf. Arrows indicate ectopic vascular sprouts. **G.** Representative brightfield images of WT control versus *dync1li1^y151^* mutants, demonstrating spinal curvature and small eyes in the mutants (black arrows). Statistics for panel B were calculated using an unpaired t-test with Welch’s correction. Data are presented as the mean ± S.D. Scale bars: 50 um.

To confirm the identity of the mutated gene, we first carried out an F0 “Crispant” experiment to confirm that transient CRISPR/Cas12a-mediated cutting of *dync1li1* at exon 12 leads to aberrant angiogenic sprouting phenotypes (**Supp. Fig. 2A**). *Tg(fli:eGFP)^y1/y1^*animals were injected with guide RNA (gRNA) and Cas12a at the one-cell stage in order to mosaically cut the endogenous *dync1li1* gene (41). Animals in which the gRNA/Cas12a efficiently cut the WT *dync1li1* gene (confirmed by PCR and fragment analysis of the resulting amplicon, **Supp. Fig. 2B-E**) displayed ectopic vascular sprouting of the ISVs, consistent with phenotypes noted in *dync1li1^y151/y151^* mutants. Animals with poor gRNA/Cas12a efficiency—leaving significant WT *dync1li1* intact—looked phenotypically normal (**Supp. Fig. 2B,E**). The presence and penetrance of phenotypes increased as the concentration of gRNA increased (**Supp. Fig. 2F**), suggesting an effect specific to Lic1 loss of function.

### LIC1-deficient ECs have abnormal vesicular trafficking consistent with inhibition of lysosomal targeting

To test the effects of LIC1 loss-of-function in human endothelial cells, we used human umbilical vein endothelial cells (HUVECs) in 3D *in vitro* collagen type 1 gel motility/sprouting assays to visualize angiogenesis *in vitro* (**Fig. 2A)** (42, 43). HUVECs pre-treated with siRNA targeting *DYNC1LI1* (siDYNC1LI1) had significantly more sprouting cells compared to siControl**-**treated cells (**Fig. 2B-D**), suggesting cross-species conservation of the role of LIC1 in limiting EC motility. Western blots confirmed siDYNC1LI1 depletion of LIC1 protein (**Fig. 2E**).

**Figure 2:**
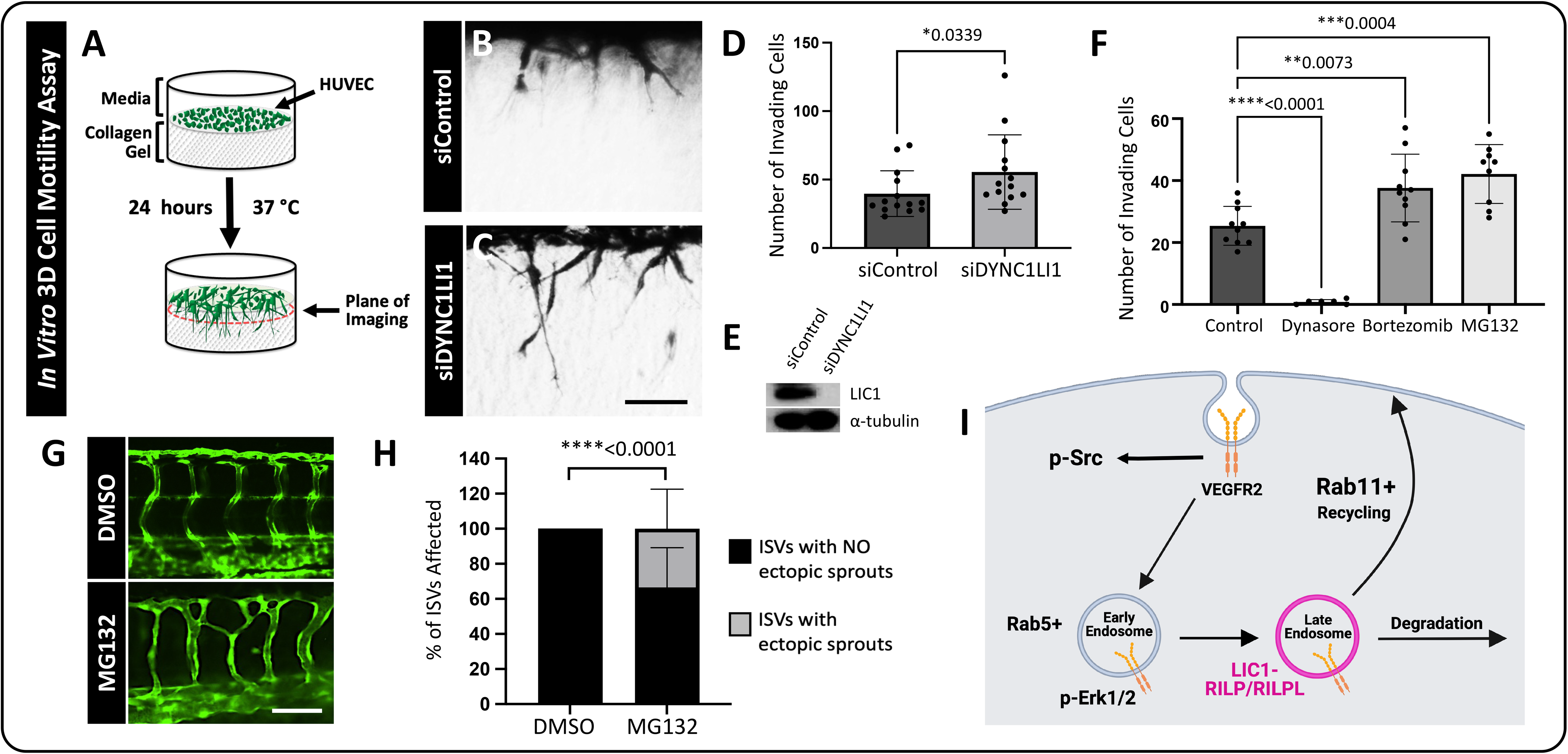
Impaired proteasomal/lysosomal function leads to increased angiogenesis *in vitro* and *in vivo*. **A.** Schematic of the *in vitro* 3D collagen type I invasion/angiogenesis assay. HUVECs are transfected with either *DYNC1LI1* or Control siRNA and seeded onto a collagen type 1 gel. After 24 hours, assays are observed for cellular invasion from the monolayer. **B, C.** Representative cross-sectional images of 3D collagen invasion assays, showing an increase in the number of invading cells in siDYNC1LI1-transfected HUVECs (B) compared to siControl-transfected cells (C). **D.** Quantification of the average number of invading cells in siControl versus siDYNC1LI1 conditions. **E**. Western blot analysis shows decreased expression of LIC1 in siDYNC1LI1-treated HUVECs compared to siControl HUVECs. Alpha-Tubulin is shown as a loading control. **F.** Quantification of the average number of invading HUVECs in 3D collagen gel assays. Assays were treated with inhibitors that block discrete stages of endocytosis. Dynasore (10uM), an early endosomal inhibitor, inhibited HUVEC invasion into the collagen gels. Bortezomib (100nM) and MG132 (10uM), which are late endosome/lysosome inhibitors, increased HUVEC invasion, consistent with siDYNC1LI1 results. **G.** Confocal images of 96 hpf zebrafish embryos treated with MG132 (10nM) or DMSO vehicle control starting at 4 hpf to investigate the effects of late endosome/lysosome inhibition on angiogenesis *in vivo*. WT *Tg(fli:eGFP)* zebrafish were used for these studies. MG132-treated fish phenocopy the *dync1li1^y151/y151^* mutant over-branching phenotype. **H.** Percentage of ISVs displaying ectopic sprouts in DMSO- or MG132-treated embryos at 96 hpf. n=20-33 embryos. **I.** Schematic of LIC1’s proposed role in regulation of endocytosis. For panels D and F, each dot represents an individual 3D collagen assay. Statistics for panel D were calculated using a Mann-Whitney test. Statistics for panel F were calculated using one-way ANOVA with Dunnett’s multiple comparisons test; omnibus ANOVA *P*-value for F (prior to the *post hoc* tests) is <0.0001. Statistics for panel H were calculated using an unpaired t-test with Welch’s correction. Data are presented as the mean ± S.D. Scale bars: 100um (B) and 50um (G).

As LIC1 is implicated in intracellular vesicular trafficking, we carried out experiments to examine how interfering with different stages of intracellular trafficking affects angiogenic sprouting. Using our 3D *in vitro* collagen type 1 gel assays with HUVECs, we treated the cultures with either the endocytosis inhibitor Dynasore, or the proteasome inhibitors MG132 and Bortezomib, which indirectly inhibit lysosomal targeting of signaling receptors through depletion of free ubiquitin (44–49). Dynasore-treated HUVECs were completely unable to invade the collagen type I gels (**Fig. 2A,F**). This observation suggests that—while endocytosis is required for angiogenic signaling (34, 35, 50, 51)**—**the distinct pro-angiogenic effect of LIC1 depletion is likely due to its involvement in a post-endocytic step. In contrast, the proteasome/lysosome inhibitors MG132 and Bortezomib stimulated HUVEC sprouting, phenocopying the response shown by siDYNC1LI1 knockdown (**Fig. 2D,F**). MG132 treatment of zebrafish also led to aberrant, excessive angiogenesis (**Fig. 2G,H**), consistent with our *in vitro* findings. Together, these data suggest that LIC1 likely exerts its effects through regulation of targeting to late endosomes or lysosomes (**Fig. 2I**).

We next sought to determine how lysosomal disruption affects vesicle distribution in endothelial cells. HUVECs were treated with the proteasome/lysosome inhibitors MG132 and Bortezomib for 1 hour, followed by labeling with LysoTracker-Deep Red for 1 additional hour. Bortezomib and MG132-treated HUVECs were found to accumulate more LysoTracker in a perinuclear cloud compared to vehicle-treated control HUVECs (**Supp. Fig. 3A,B**). In addition, MG132- and Bortezomib-treated HUVECs exhibited significantly higher levels of Rab11a+ recycling endosomes in the perinuclear and plasma membrane regions of cells compared to the vehicle-treated controls (**Supp. Fig. 3C,D**). The increased accumulation of LysoTracker, accompanied by the increase in Rab11a+ endosomes, suggests that pro-angiogenic signals might not be properly extinguished in cells with impaired lysosomal targeting. Instead, signaling receptors may undergo recycling to the plasma membrane, or continue to accumulate and persist signaling, thereby driving excess angiogenesis.

### LIC1-deficient ECs have increased surface VEGFR2 levels and p-SRC activation, coupled with decreased VE-Cadherin levels

The *in vivo* findings detailed above indicate that loss of LIC1 and loss of lysosomal targeting led to similar angiogenic sprouting phenotypes. Since VEGFR2 is known to be a driver of angiogenesis and is also known to be regulated by trafficking to the lysosome (20, 35, 38, 52), we next sought to determine whether LIC1 deficiency impacts VEGFR2 localization in HUVECs. HUVECs were treated with either siDYNC1LI1 or siControl and stimulated with the VEGFR2 ligand VEGF-A for 1 hour. Cells were then fixed and VEGFR2 localization was examined by immunofluorescence microscopy, with and without permeabilization, to determine total and surface VEGFR2 levels, respectively (**Fig. 3**). While there was no difference in the total VEGFR2 protein levels between siControl and siDYNC1LI1 cells (membrane permeabilization with cell fixation, **Fig 3A,B**), siDYNC1LI1 cells had higher levels of VEGFR2 on the cell surface compared to siControl HUVECs (no membrane permeabilization with cell fixation, **Fig. 3A,C**), an effect previously shown to promote excess angiogenesis (53).

**Figure 3:**
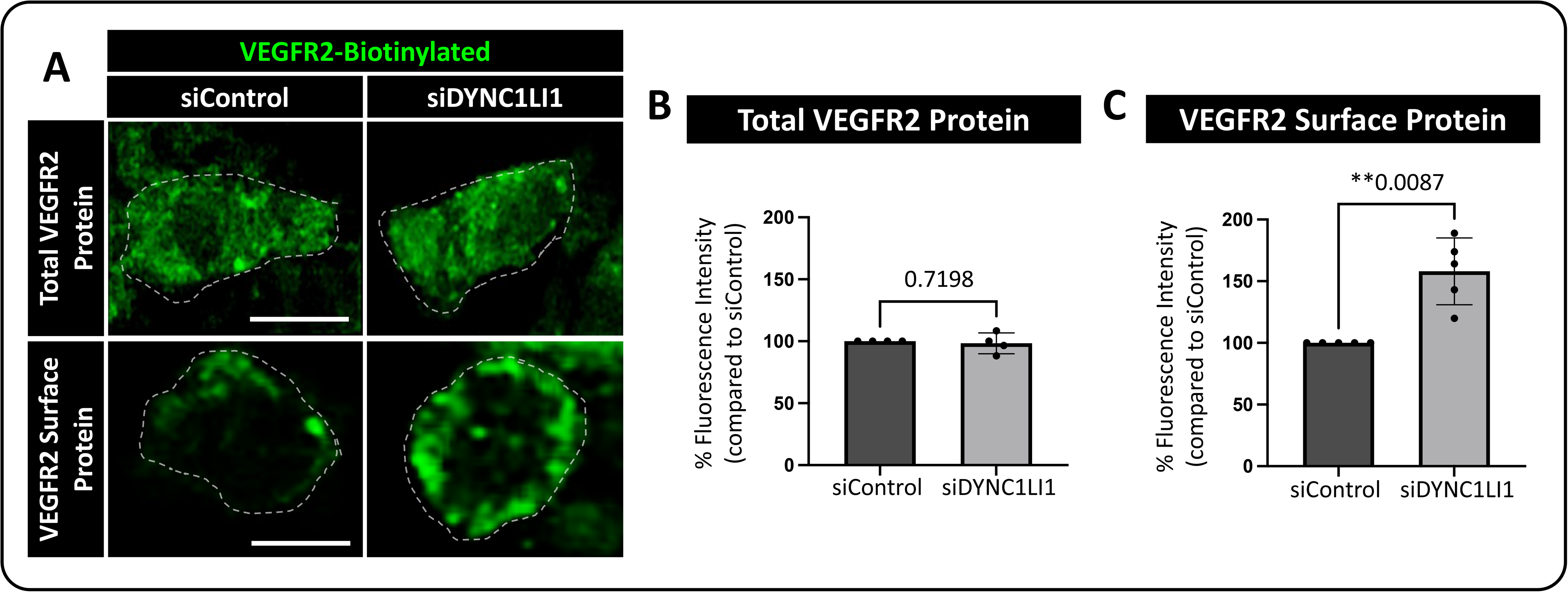
VEGFR2 protein surface localization is increased in LIC1-deficient HUVECs. **A.** Representative single plane images of total VEGFR2 versus surface expression in HUVECs treated with siDYNC1LI1 versus siControl. Cells were simulated with 40 ng/ml VEGF-A for 1 hour then fixed and immunostained. Dashed lines highlight individual cell borders. **B**. Quantification of total (whole cell, permeabilized) VEGFR2 protein levels. **C.** Quantification of surface (non-permeabilized) VEGFR2 protein levels. For panels B and C, each dot represents data from an independent experiment. Statistics for panels B and C were calculated using an unpaired t-test with Welch’s correction. Data are presented as the mean ± S.D. Scale bars: 10um.

Phosphorylation of ERK1/2 and SRC occurs downstream of VEGFR2 signaling (36, 54). To determine whether these pathways were affected by loss of LIC1, we compared activation after VEGF-A treatment in siDYNC1LI1 or siControl HUVECs. No overt differences were noted in p-ERK1/2 activation between siControl and siDYNC1LI1 HUVECs, with or without VEGF-A stimulation (**Fig. 4A**). However, siDYNC1LI1 HUVECs had higher baseline levels of p-SRC than siControl HUVECs (no VEGF-A stimulation), which modestly increased upon VEGF-A stimulation (**Fig. 4A**). Treatment of HUVECs with the endocytosis inhibitor Dynasore led to nearly complete suppression of p-ERK1/2—consistent with published reports stating that VEGFR2 must be endocytosed to activate ERK (31, 50, 55)—yet showed high p-SRC activation even at baseline (no VEGF-A stimulation; **Fig. 4B**). Finally, HUVECs treated with MG132 or Bortezomib showed comparable p-ERK1/2 activation but increased p-SRC activation relative to vehicle-treated controls (**Fig. 4C**). Together, our data suggest that LIC1-deficiency and inhibition of lysosomal targeting results in sustained activation of SRC in the presence or absence of VEGF-A stimulation. This prolonged SRC activation may drive excess angiogenesis in siDYNC1LI1 HUVECs and *dync1li1^y151/y151^* mutant zebrafish. Additionally, the sustained p-SRC activity observed in Dynasore-treated HUVECs suggests that VEGFR2 is able to activate SRC from the plasma membrane to drive increased angiogenesis in the absence of VEGFR2 endocytosis.

**Figure 4:**
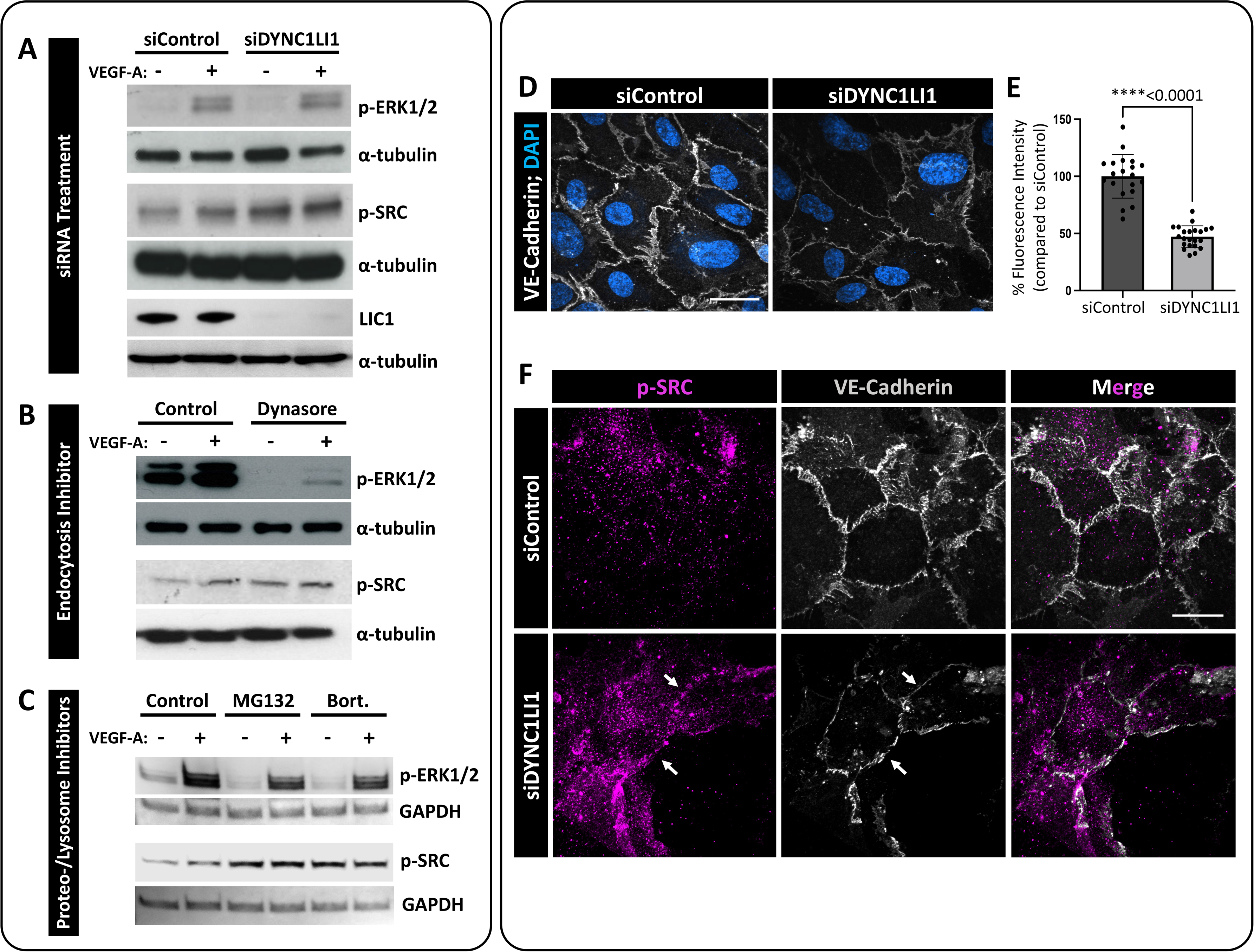
p-SRC activation is increased in LIC1-deficient HUVECs. **A-C.** Representative Western blots of p-SRC and p-ERK1/2 activation in response to VEGF-A stimulation in siDYNC1LI1-transfected HUVECs compared to siControl cells (A); Dynasore-treated HUVECs compared to DMSO vehicle control treated cells (control, B); and HUVECs treated with MG132 or Bortezomib compared to DMSO vehicle control treated cells (control, C). GAPDH or a-tubulin are shown as protein loading controls. LIC1 levels following siRNA treatment is shown to confirm protein suppression (A). **D.** Immunofluorescent labeling of VE-Cadherin (grey) and nuclei (blue) in HUVECs transfected with siDYNC1LI1 or siControl. **E.** Quantification of VE-Cadherin levels. Statistics for panel E were calculated using an unpaired t-test with Welch’s correction. Data are presented as the mean ± S.D. **F.** Immunofluorescent labeling of VE-Cadherin (grey) and p-SRC (magenta) in HUVECs transfected with siDYNC1LI1 or siControl. White arrows highlight EC-EC boundaries. Scale bars: 5um.

Since p-SRC has been shown to be involved in cellular motility, vascular permeability, adhesion, and proliferation (56–58), we wanted to determine if the abnormal SRC activation noted in siDYNC1LI1 ECs altered p-SRC localization and EC-EC junction formation. SRC-mediated phosphorylation of VE-Cadherin can lead to decreased junction formation and vessel wall stabilization, perhaps accounting for the increased motility noted in our siDYNC1LI1-treated ECs (51, 59). Immunolabeling of VE-Cadherin in siControl-versus siDYNC1LI1-treated HUVECs showed that VE-Cadherin levels were decreased approximately 50% in response to LIC1 suppression (**Fig. 4D,E**). The loss of VE-Cadherin levels correlated with increased p-SRC accumulation at EC-EC boundaries in the siDYNC1LI1 HUVECs (**Fig. 4F**), suggesting that increased and sustained p-SRC activation in siDYNC1LI1 HUVECs helps maintain a loose EC-EC junction to allow for continued vessel outgrowth.

### Increased Rab11a-mediated endosomal recycling drives excess angiogenesis

Rab binding proteins are key regulators of different steps of intracellular vesicular transport but it remains unclear how these proteins are required for angiogenesis. To examine the effects of loss of LIC1 on different Rab proteins, we activated VEGFR2 signaling in siDYNC1LI1 or siControl HUVECs by VEGF-A treatment and then fixed and immunostained the cells for Rab7 (late endosomes/lysosomes) or Rab11a (recycling endosomes). We observed slightly elevated Rab7 and increased Rab11a staining in siDYNC1LI1 HUVECs compared to siControl HUVECs (**Fig. 5A-D**). Western blot analysis confirmed the increased Rab11a protein levels in siDYNC1LI1 HUVECs (**Fig. 5E**), while qPCR analysis of siControl and siDYNC1LI1 HUVECs revealed minimal changes in transcript levels of either Rab7a or Rab11a (**Fig. 5F**). Together, these data suggest that loss of LIC1 may lead to an increase in Rab11a-driven recycling of VEGFR2 to the plasma membrane to promote angiogenic sprouting.

**Figure 5:**
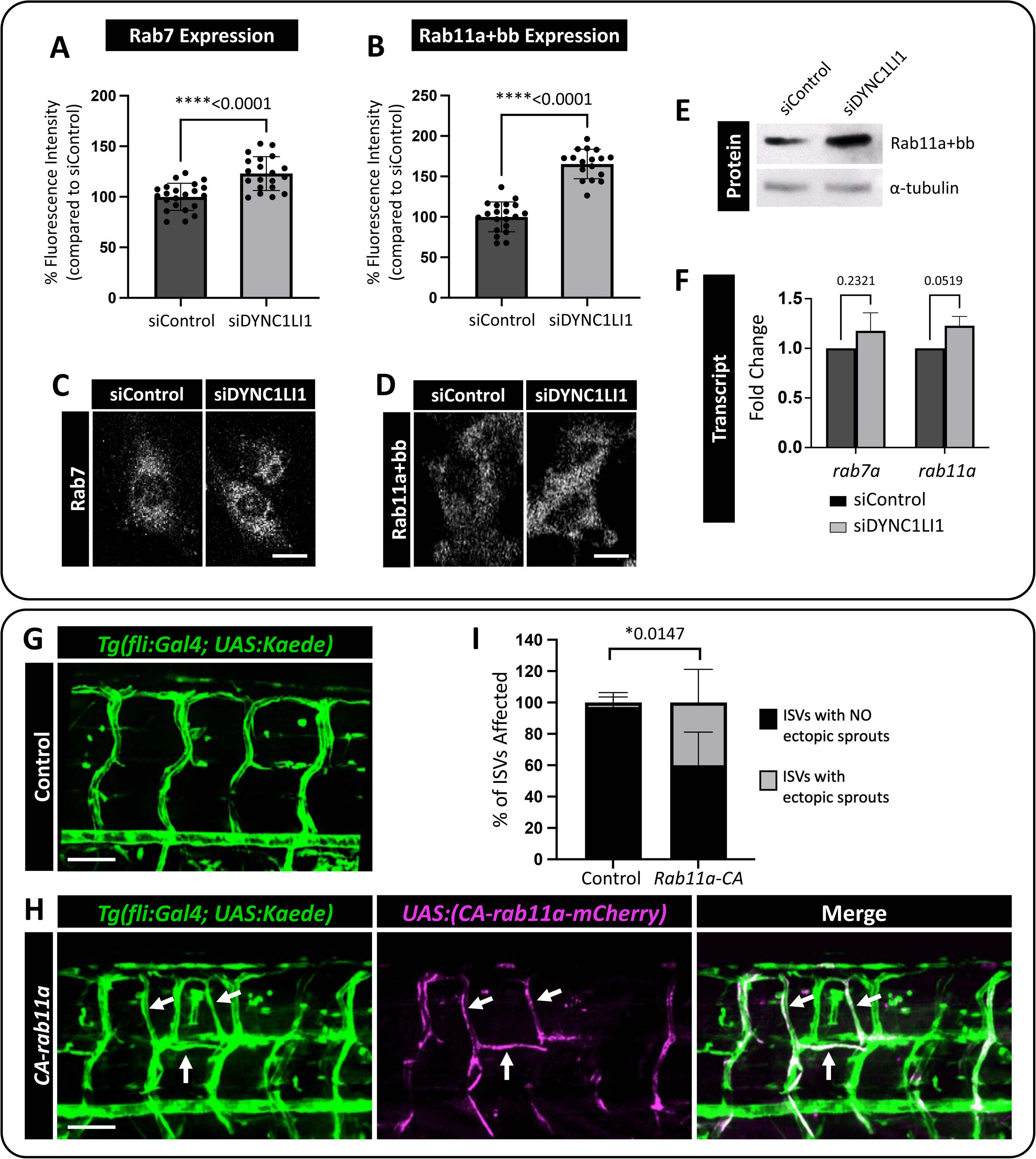
Constitutive Rab11a activation leads to ectopic angiogenic sprouting. **A,B.** Quantification of Rab7 (A) and Rab11a+bb (B) positive vesicles in siDYNC1LI1 HUVECs versus siControl-treated cells. **C,D.** Representative images of HUVECs immunostained for Rab7 (C) and Rab11a+bb (D) following siControl or siDYNC1LI1 transfection and treatment with 40 ng/mL VEGF-A for 10 min. **E.** Western blot analysis shows increased expression of Rab11 in siDYNC1LI1 HUVECs compared to siControl cells. **F.** qPCR reveals no change in the transcript levels of either Rab7 or Rab11 in siDYNC1LI1 cells, suggesting that the increased protein levels of Rab11 occur post-transcriptionally. **G,H.** Confocal images of *Tg(fli:Gal4); Tg(UAS:Kaede)* zebrafish embryos injected with a Tol2-integratable *UAS:CA-rab11a-mCherry* DNA plasmid to express constitutively active Rab11a protein in the endothelium in a mosaic fashion (H) versus carrier injected controls (G). Blood vessels are shown in green; mosaic expression of *UAS:CA-rab11a-mCherry* is shown in magenta. Endothelial cells expressing *UAS:CA-rab11a-mCherry* and *Tg(fli:Gal4); Tg(UAS:Kaede)* appear white when the images are merged. Sites of *UAS:CA-rab11a-mCherry* expression (white arrows) phenocopy the increased angiogenesis noted in *dync1li1^y151/y151^* mutants. **I.** Percentage of ISVs displaying ectopic sprouts in carrier injected controls versus *UAS:CA-rab11a-mCherry-*injected embryos at 96 hpf. n=5 embryos. For panels A,B, each dot represents an individual cell. Panels A,B are representative of three independent experiments and statistics were calculated using an unpaired t-tests. For panel F, n=3. Statistics for panel F were calculated using one-sample t-tests to compare siDYNC1LI1 fold change values to hypothetical value 1. Data are presented as the mean ± S.D. Statistics for panel I were calculated using an unpaired t-test with Welch’s correction. Data are presented as the mean ± S.D. Scale bars: 10um (C,D) and 50uM (G,H).

To test the hypothesis that increased Rab11a activity can alter angiogenic potential during zebrafish vascular development *in vivo*, we expressed constitutively-active Rab11a (CA*-rab11a*) specifically in ECs using the UAS/gal4 system (60). Tol2(*uas:CA-rab11a-mCherry)* DNA was injected into *Tg(fli:gal4); Tg(uas:Kaede)* double transgenic embryos at the one-cell stage, and mosaic CA*-rab11a* expression was tracked at subsequent stages using the mCherry reporter. Embryos with mosaic patches of vascular mCherry expression were imaged at 96 hpf (**Fig. 5G,H**). Ectopic angiogenic sprouts were more frequently found to be mCherry-positive (highlighted by white arrows; **Fig. 5G-I**) than mCherry-negative, supporting the hypothesis that increased Rab11a activity leads to a cell-autonomous increase in angiogenesis. This further suggests that the *dync1li1^y151/y151^* mutants might have enhanced Rab11a-mediated endosomal recycling and suppressed lysosomal targeting.

### *rilpl1/rilpl2* mutants phenocopy *dync1li1^y151^* mutant phenotypes

Our results suggest that increasing recycling of endosomes via increased Rab11a activity promotes excess angiogenesis. To further examine whether decreased Lic1/Rab-dependent lysosomal degradation promotes angiogenesis, we carried out additional studies on the zebrafish *rab-interacting lysosomal protein like 1*/*2 (rilpl1/2)* genes. Previous work has shown that the LIC proteins comprise an N-terminal GTPase-like domain that binds to the dynein heavy chain and a C-terminal adaptor-binding domain that binds to various Rab effectors (9, 61–64). LICs can thus link the dynein motor complex to Rabs to regulate endocytic trafficking and promote the transition from late endosome to lysosome (**Fig. 6A-C**) (29, 62, 64, 65). The Lic1 mutation in the *dync1li1^y151^* zebrafish removes the last 36 amino-acid residues from the C-terminal domain of the WT protein and introduces 3 amino-acid residues including a stop codon in frame (**Supp. Fig. 1A, Fig. 6B**). To examine whether this truncated protein has reduced binding affinity for RILP, we generated GST-fused C-terminus of WT and mutant human LIC1 proteins (i.e. insertion of the 19bp sequence and removal of exon 13, **Supp. Fig. 1A, Fig. 6B,C**) for GST pulldown assays. We observed that WT LIC1-GST (GST-WT LIC1 CT) bound robustly to GFP-RILP (anti-GFP, RILP), whereas mutant LIC1-GST (GST-MUT LIC1 CT) had markedly reduced binding to GFP-RILP (anti-GFP, RILP) (**Fig. 6D,E**).

**Figure 6:**
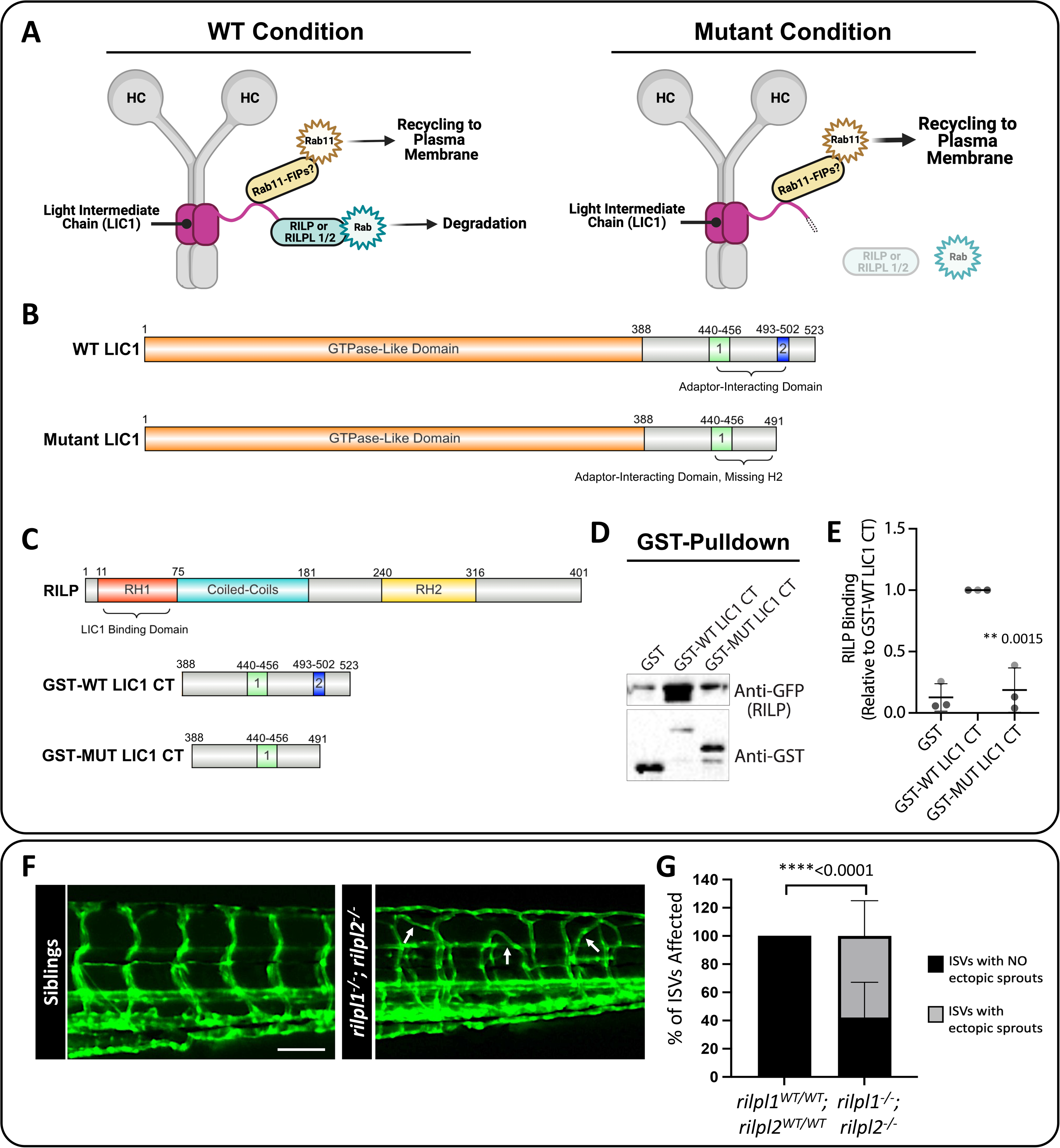
*rilpl1/2* mutants recapitulate the increased angiogenesis phenotype seen in *dync1li1^y151/y151^* mutants. **A.** Schematic representation of the role of RILP or RILPL1/2 in bridging LIC1 and Rab binding. RILP is an adaptor for Rab7, and RILPL1/2 are adaptors for Rabs 8, 10, and 36, all of which regulate lysosomal biogenesis and degradation. Mutations to RILP or RILPL1/2 are predicted to impair Rab binding to endosomes and decreased lysosomal degradation. **B.** WT LIC1 domains: GTPase Like Domain (1–388) and Adaptor-Interacting Domain (Helix 1 (440–456) and Helix 2 (493–502)). Mutant LIC1 domains: GTPase Like Domain (1-388) and Adaptor-Interacting Domain (Helix 1 (440-456), Helix 2 is deleted in this mutant). The mutant LIC1 introduces a 19 bp insertion at the end of exon 12. This insertion adds 2 amino acids: aspartic acid (Asp), leucine (Leu), and a premature “STOP” codon resulting in the complete loss of exon 13 and the H2 domain. **C.** Schematics of the proteins used for GST-binding assays. RILP: RILP Homology 1 (RH1) (11-75), Coiled-Coils (75-181), and RILP Homology 1 (RH2) (240-316) domains. RILP’s RH1 domain binds to the LIC1 Adaptor-Interacting Domain. GST-WT LIC1 CT (GST-fused wild type human LIC1 c-terminal adaptor-interacting domain, a.a. 388-523). GST-MUT LIC1 CT (GST-fused mutant human LIC1 c-terminal adaptor-interacting domain a.a. 388-491, including the introduced Asp, Leu, STOP found in the zebrafish mutant, which is predicted to change protein conformation). Schematics created with Ibs.renlab.org. **D.** GST pulldown of WT human LIC1 C-terminus (GST-WT LIC1 CT) or the C-terminus bearing the equivalent mutation found in the *dync1li1^y151^* mutant zebrafish (GST-MUT LIC1 CT). Purified RILP protein displays decreased binding to the mutant LIC1 C-terminus compared to WT LIC1 C-terminus. **E.** Quantification of the Western blot data assessing RILP binding to GST-WT LIC1 CT versus GST-Mut LIC1 CT. n=3. **F.** Confocal images of *rilpl1/2* double-mutants at 96 hpf. **G.** Percentage of ISVs displaying ectopic sprouts in *rilpl1/2* double-mutants compared to WT control siblings at 96 hpf. n=20 embryos. Statistics were calculated using an unpaired t-test with Welch’s correction. Data are presented as the mean ± S.D. Schematics created with BioRender.com. Scale bars: 50um.

To further test the functional consequence of disrupted interaction between Lic1 and Rilpl proteins *in vivo*, we generated *rilpl1^stl842^/ rilpl2^stl843^* double mutant zebrafish using CRISPR/Cas9 genome editing (**Supp. Fig. 1B,C**) (28). Analysis of *rilpl1^stl842/stl842^/ rilpl2^stl843/stl843^*double homozygous mutants versus control siblings at 96 hpf shows aberrant angiogenesis, phenocopying the *dync1li1^y151/y151^* mutants (**Fig. 6F,G**). Taken together, these data support the conclusion that the *dync1li1^y151/y151^* mutation disrupts Rilpl1/2 binding, to decrease lysosomal degradation and, consequently, increase recycling, increase cell-surface VEGFR2, and promote excess angiogenesis (**Fig. 7**).

**Figure 7:**
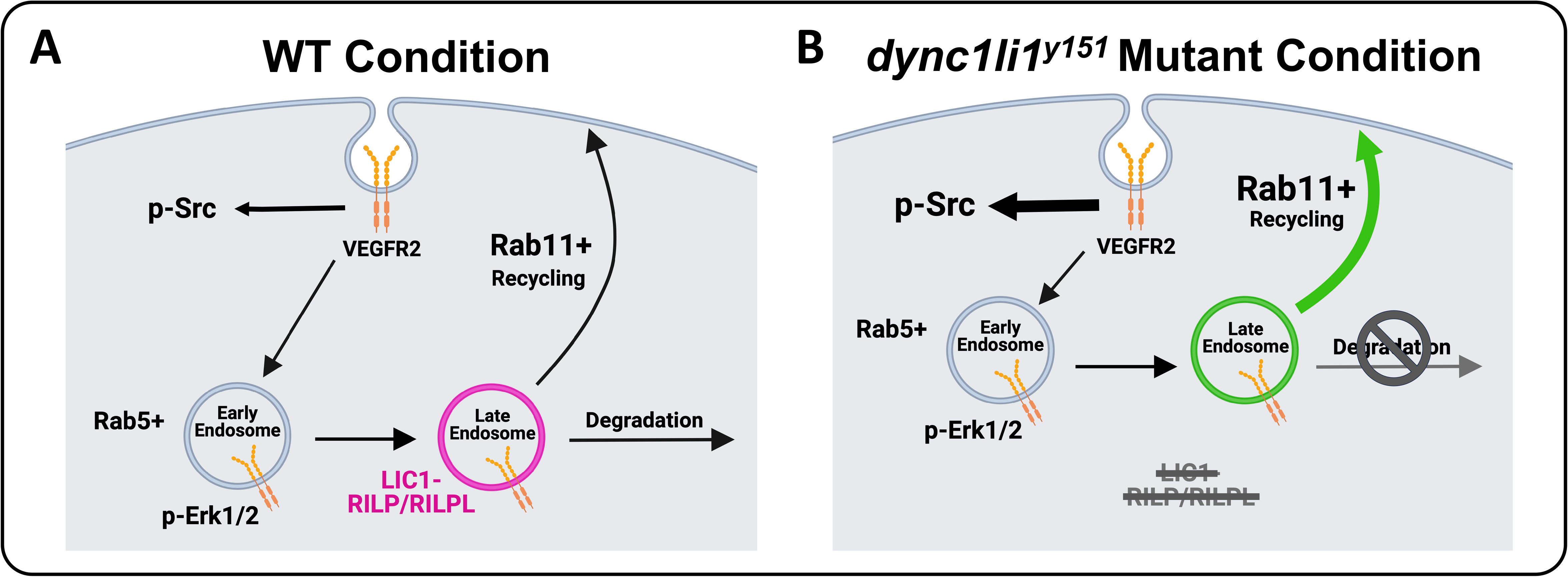
A model of the role of LIC1 in angiogenesis. **A.** LIC1 regulates binding of Rab adaptor proteins—in particular RILP and RILPL1/2—to augment endosomal recycling versus degradation. **B.** In the absence of the LIC1 C-terminal domain, RILP and RILPL1/2 are unable to bind LIC1 efficiently, leading to increased Rab11-mediated endosomal recycling and decreased Rab7-mediated endosomal degradation. Increased surface expression of VEGFR2 protein leads to overactivation of p-SRC and increased endothelial cell motility. Schematics created with BioRender.com.

## DISCUSSION

### The role of LIC1-regulated vesicular trafficking in angiogenesis

The regulation and trafficking of cargo by the dynein motor complex is central to many different cellular processes, including endocytosis (66). While studies have shown that mutations in the dynein motor machinery are often sufficient to cause motor neuron diseases (i.e. Huntington Disease, Amyotrophic Lateral Sclerosis, and Alzheimer’s Disease, among others (15, 23, 26, 67, 68)), until this study, the role of LIC1 in endothelial cell biology and angiogenesis had remained unknown. Here, we identify and characterize a novel zebrafish mutant in the *dync1li1* gene, *dync1li1^y151^*, that exhibits increased angiogenesis in multiple vascular beds—including the ISVs and sub-intestinal vascular plexus (**Fig 1C-F**). While angiogenesis is critical for vascular expansion during development and tissue repair during wound healing, abnormal angiogenesis has been shown to contribute to variety of diseases/disorders including cancer, diabetes, stroke, arthritis, and psoriasis (39, 69–71). Thus, discovering new mechanisms to promote or dampen blood vessel growth will continue to have impacts on the development of targeted pro- and anti-angiogenic therapies.

The structure of LIC1 allows it to simultaneously interact with the dynein motor complex and cargo adaptors at its C-terminus (63). Previous work has shown that LIC1 colocalizes with the adaptors RILP or RILPL1/2 primarily on late endosomes (72), a point that might speak to the critical role of LIC1 in regulating lysosomal degradation. Our functional findings *in vivo* suggest that there is a unique role for the LIC1-RILPL1/2 interaction in regulating vesicles at the late endosome/lysosome transition. The exact intracellular cues that promote the LIC1-RILPL1/2 interaction remain unclear. However, from a structural standpoint, prior research suggests that the small, complex binding domains between the C-terminal of LIC1 and RILP or RILPL1/2 facilitate the specificity of the interaction (61, 72, 73). The zebrafish *dync1li1* mutation we identified supports this finding. The identified mutation at the *dync1li1* exon12/13 boundary likely results in a truncated Lic1 without its C-terminal domain, and mutations in *rilpl1 ^stl842^/ rilpl2^stl843^*in the zebrafish phenocopy the *dync1li1^y151/y151^* mutant. Together, these data support the conclusion that deficient LIC1-RILPL1/2 interactions can drive excessive angiogenesis.

Due to the role of LIC1 in regulating endocytic maturation, we investigated the effects of endocytic compartment maturation on vessel sprouting. *In vitro*, HUVECs were treated with endocytosis inhibitors Dynasore versus the proteasomal/lysosomal targeting inhibitors MG132 or Bortezomib (74, 75). MG132 has been shown to “trap” both randomly moving lysosomes and lysosomes surrounding the centrosome to decrease lysosomal function (76), while Bortezomib blocks autophagic flux and vesicular degradation activity, without interfering with autophagosome and lysosome fusion (47, 77). Interestingly, both can inhibit lysosomal targeting of signaling receptors by depleting pools of free ubiquitin, which could be the mechanism underlying our phenotype (44–49). Importantly, while Dynasore-treated HUVECs do not invade into collagen type I gels, MG132- and Bortezomib-treated HUVECs have a significantly higher number of invading cells compared to control (**Fig. 2F**)—similar to the findings seen with siDYNC1LI1 HUVECs (**Fig. 2D**)—suggesting increased signaling activity under these conditions. *In vivo*, zebrafish embryos treated with MG132 showed increased angiogenic sprouting along the ISVs, consistent with *dync1li1^y151/y151^*mutants (**Fig. 2G,H**). These findings suggest that the *dync1li1* mutation likely disrupts the late endosome-lysosome transition but does not affect the regulation of early endosomes or the initial stages of endocytosis. As a consequence, Rab11a-mediated vesicular recycling is predicted to be increased in *dync1li1^y151/y151^* mutants and siDYNC1LI1 HUVECs, promoting the cell surface availability of pro-angiogenic receptors such as VEGFR2 (**Fig. 3-4**). These findings are consistent with previous studies demonstrating that an increase in the number of late endosomes, without induction of lysosomal proteolysis, can result in the retention of signaling cargos (72, 78–80). Furthermore, it was shown that aggressive cancer cells can exhibit increased accumulation of lysosomes near the plasma membrane, prior to lysosomal exocytosis, resulting in increased cellular proliferation and tumor invasion (78, 81). Together, these data suggest that disruption of lysosomal targeting can lead to the overexpression of VEGFR2 and other pro-angiogenic receptors at the plasma membrane to promote angiogenesis.

### Enhanced pro-angiogenic signaling correlates with increased vesicular recycling

VEGFR2 transduces intracellular signals through a variety of mediators including the SRC and PLC/PKC/ERK1/2 pathways. These signaling pathways are responsible for regulating vascular permeability, cell survival, migration, proliferation, differentiation, invasion, and angiogenesis (82–85). Previous work has shown that VEGFR2 must undergo endocytosis for the initiation of pro-angiogenic signaling via ERK1/2 (31, 50, 55); however, the necessity of endocytosis for phosphorylation and activation of SRC has been less clear.

Interestingly, we see unchanged p-ERK1/2 activity in our models (siRNA suppression of *DYNC1LI1* or MG132/Bortezomib treatment). However, p-SRC, which we suspect can be activated from the cell surface in an endocytosis-independent manner, is increased in accordance with inhibition of late endosome/lysosomal maturation and following siRNA suppression of DYNC1LI1 (**Fig. 4**). This suggests that amplified p-SRC activation is contributing to the increased sprouting. We also demonstrated that VE-Cadherin levels decrease while p-SRC accumulates at EC-EC boundaries in siDYNC1LI1 HUVECs (**Fig. 4**). Together, these data support a mechanism by which endothelial cells loosen their junctions to allow for increased angiogenic sprouting in accordance with increased surface availability of pro-angiogenic receptors, such as VEGFR2 (**Fig. 3,4**). While we use VEGFR2 as an example in this study, there are a host of other signaling cargoes that could be modified via this process, and additional work will need to be done to define the causative versus correlative role of VEGFR2 in contributing to the angiogenic sprouting phenotypes noted. Despite this, these studies define a central role for LIC1 in regulating angiogenesis via vesicular recycling.

Using constitutively active Rab11a (CA*-rab11a*), we show how altering EC-autonomous vesicular recycling can regulate angiogenesis *in vivo* on a wild-type background (**Fig. 5**). As Rab11a is responsible for recycling cargoes back to the plasma membrane, this can allow for sustained availability of pro-angiogenic signals without the need to synthesize de novo protein. In fact, in other contexts, overexpression of Rab11a has been shown to contribute to the proliferation and invasion of cancer cells via increased signaling of ERK, PI3K/AKT, EGFR, WNT, and MMP2, consistent with our findings that increased Rab11 activity can drive angiogenesis (86). Further, *rilpl1^stl842^/ rilpl2^stl843^* mutant zebrafish also exhibit an ectopic sprouting phenotype (**Fig. 6**). By using these *rilpl1^stl842^/ rilpl2^stl843^*mutant zebrafish, we show how potential disruption of the Lic1-Rilpl1/2-Rab protein complex can result in the suppression of lysosomal degradation leading to an increase in ectopic vascular sprouting. While intriguing, further work will need to be done to understand the long-term consequences of impaired lysosomal targeting on developmental processes, as all the mutant alleles described here are lethal by 2 weeks post-fertilization.

In summary, we demonstrate via the characterization of novel zebrafish mutants, *dync1li1^y151^* and *rilpl1^stl842^/ rilpl2^stl843^*, and through *in vitro* assays using HUVECs, that impaired maturation of endosomes and/or targeting to lysosomes of pro-angiogenic signaling receptors—such as VEGFR2—leads to enhanced or sustained angiogenesis. In the *dync1li1^y151^*and *rilpl1^stl842^/ rilpl2^stl843^* mutants, and in the siDYNC1LI1-treated HUVECs, this is likely driven by a decrease or halt in lysosomal targeting due to a loss of the Lic1-Rilpl1/2 interaction. Based on these findings, we place LIC1 as a key mediator in the regulation and/or maturation of endosomes via differential interactions with Rab adaptor proteins, ultimately resulting in targeted regulation of cellular motility and angiogenesis (**Fig. 7**).

## METHODS

### Zebrafish transgenic lines and husbandry

Zebrafish (*Danio rerio*) *dync1li1^y151^* mutants were isolated during an ENU mutagenesis screen. New lines for this study were generated using CRISPR/Cas9 mutagenesis, injecting the guide RNAs at 150 pg/nl per embryo (87): *rilpl1^stl842^* and *rilpl2^stl843^*. Embryos were raised and maintained within the *Zebrafish Consortiums* at the National Institutes of Health and Washington University School of Medicine. Zebrafish husbandry and research protocols were reviewed and approved by the NIH/NICHD and Washington University in St. Louis Animal Care and Use Committees.

### Zebrafish Crispant generation, genotyping, and microscopy

Fluorescent images of 72-120 hpf zebrafish were collected using the W1 Spinning Disk confocal microscope, a Fusion camera, and the Nikon Eclipse Ti2-E base with a 20x objective. Embryos were anesthetized prior to imaging using tricaine methanesulfonate (MS-222; Western Chemical, Inc; NC0872873) and embedded in low melting point agarose (IBI Scientific; #IB70056) on a 35-mm glass-bottom petri dish (MatTek; #P35G-1.5-20-C) for imaging. Blinded analysis was performed on all images. Embryos were then genotyped using pre-designed primers (see below for PCR protocol and primers).

Confocal microscopy was performed in order to compare the onset of vascular defects of homozygous mutants versus sibling controls (homozygous WT). The intersegmental vessels (ISVs) and sub-intestinal vessels (SIVs) were chosen as vascular areas for comparison due to minimal developmental variation. To quantify vascular defects over time, the percentage of ISVs displaying ectopic sprouts and the average number of ectopic sprouts per ISV were quantified at different time points (10 ISVs analyzed per zebrafish).

CRISPR mutant generation and Genotyping:

Zebrafish were genotyped using the following primers:

#### dync1li1^y151^

Common FW: TGTCATTTCTGGTGGCAGGA

Mutant RV: GCTCACCGTTGTTGATGTCA

WT RV: GATCGCTCACCTGACTTGC

#### _rilpl1_^stl842^

FW: TGTAAAACGACGGCCAGTGCATTACCTCCTGTAGTAG

RV: GTGTCTTGACCACCAGGAGGCGCTC

CRISPR Guide: GGGTGAAGCGCGGCCGGTTA

#### _rilpl2_^stl843^

FW: TGTAAAACGACGGCCAGTCCAGCGACAGGTTGTATTTGTTC

RV: GTGTCTTCGCAAGCTTTCGACAAGGACGTG

CRISPR Guide: GGGTGGAAACCGGGAGATCT

Primers were designed using the online browser Primer3web (version 4.1.0), then purchased from Integrated DNA Technologies (IDT). Online browsers Ensembl (release 110) and Primer-BLAST (NCBI) were used as genomic reference tools.

Zebrafish were fin-clipped under anesthesia using 0.1% MS-222. Genomic DNA was extracted using extraction solution (E7526; Millipore Sigma), tissue preparation solution (T3073; Millipore Sigma), and Neutralization Solution B (N3910; Millipore Sigma). PCRs were performed using DreamTaq Polymerase (ThermoFisher Scientific; K1081) with primers designed to distinguish between wild type, heterozygous, or homozygous mutant carriers. For the *dync1li1* mutant specifically: The primers were designed so the mutant reverse primer sits directly on the 19 bp insert mapped to be the *lic1^y151^* mutation site. The wildtype reverse primer was designed across the DNA site that is disrupted by the insertion in the mutant. This primer is only able to amplify the wild type allele. A common forward primer was designed for both alleles. The PCR reaction utilized all three primers at once, allowing amplification of either a single wildtype band, two bands that are 19bp different in size (heterozygous mutants), or a single mutant band (homozygous mutant siblings). PCR products were run and sized using an Agilent 5300 Fragment Analyzer (41).

PCR Primer Annealing Temperatures:

*dync1li1: 53 °C*
*rilp1: 56 °C*
*rilp2: 61 °C*

F0 Crispant injections were carried out on the *Tg(fli:eGFP)^y1^* transgenic line. Cas12a and the CRISPR guide RNA (GGTGGCAGGAGGTCAGACCAGTGA) were injected with a dose of 50 or 100 pg/nl guide RNA at the one cell stage with 10uM/uL Cas12a protein and allowed to develop until 96 hpf. Spinning disk confocal imaging was carried out on embryos at 96 hpf as outlined above. Cutting efficiency was assessed by PCR of gDNA extracted from individually injected and imaged embryos. PCR products were run and sized using an Agilent 5300 Fragment Analyzer (41). Cutting efficiency was determined by comparing the percent of PCR amplicons present not at the WT amplicon size to the percent WT amplicon present. Primers: FW 5’-GATGAGCAAATGATTCGGAAAT; RV 5’-AGAGCCTGCTTTCTTTGTCAAG.

### Zebrafish Injections

For all expression studies in the zebrafish: constructs were injected into one-cell stage zebrafish embryos. Confocal microscopy was used to image the effects of the injected constructs on wild type or the *dync1li1^y151^* mutant phenotype. The CA-*rab11a* construct (88) was injected at 50 ng/pl for analysis on the *Tg(fli:gal4);Tg(UAS:Kaede)* background for analysis.

### Endothelial 3D Invasion Assay

Human umbilical vein endothelial cells (HUVECs; Lonza) were cultured in 15 mg bovine hypothalamus extract (#02-102, Sigma-Aldrich), 0.05% Heparin (#H3393, Sigma-Aldrich) and 20% FBS (Gibco) in M199 base media (#11150059, Gibco) on 1 mg/ml gelatin-coated tissue culture flasks. HUVECs were used from passages 2**–**5. Endothelial 3D invasion assays were performed using collagen type I (Corning; #354249, Acid Extracted from Rat Tail) gels and HUVECs transfected with siRNAs: *DYNC1LI1* versus a non-specific negative control. HUVECs were seeded on top of the collagen gel at a density of 40,000 cells per well and allowed to invade the gel for 24 hours. Assays were fixed in 2% paraformaldehyde (PFA) and processed for further analysis. Culture media for the assays contained L-ascorbic acid (AA) (#A61-25, Fisher Scientific), FGF (#233-FB-025/CF, R&D systems), and IGF-2 (#292-G2-250, R&D systems). Additionally, FGF, SCF (#255-SC-010/CF, R&D systems), IL3 (#203-IL-010/CF, R&D systems), and SDF1a (#350-NS-010/CF, R&D systems) were supplied in the collagen gel (42, 89, 90). Images were taken with EVOS M5000.

### siRNA transfection

Validated Silencer Select small-interfering RNAs (siRNAs) targeting human *DYNC1l11* (ThermoFisher Scientific; 4390824) or non-specific negative control siRNAs (ThermoFisher Scientific; 4390847) were purchased and resuspended in RNase-free H_2_O to generate a 25μM stock concentration. siPORT Amine (Invitrogen; #AM4502) was used as the transfection reagent. HUVECs were transfected as per the siPORT Amine transfection protocol with 5 nM of siRNA on days 1 and 3, then used for downstream analysis on day 4.

### Inhibition of Endosomal Trafficking

HUVECs were treated with endosomal inhibitors, 10 uM Dynasore (Tocris Bioscience; 2897), 10 uM MG132 (Tocris Bioscience; 1748), and 100 nM Bortezomib (EMD Millipore; 504314) after culturing to a confluent monolayer in a tissue culture flask. For VEGF-A treated cells, 40ng/mL of VEGF-A was added for 1 hour post inhibition. Alternatively, inhibitors were added 30 minutes after seeding HUVECs onto collagen type 1 gels for 3D invasion assays. For endosomal trafficking inhibition in zebrafish, embryos were treated with either 15 uM MG132 (Tocris Bioscience; 1748) or DMSO (Sigma; D4540) via fish water starting at 4 hpf.

### Western blot Assays

HUVECs were lysed using RIPA buffer (Pierce; 89900) with Protease Inhibitor (Roche; 04693132001) and Phosphatase Inhibitor (Sigma; P8340). 20-30 μg protein was electrophoresed on a 10-20% SDS-PAGE gel (Invitrogen; XP10200BOX), transferred to a PVDF membrane (ThermoFisher; 88518), and then blocked in 5% BSA (phosphorylated proteins) or 5% nonfat dry milk in Tris-buffered saline with 0.1% Tween 20 (TBST) for 1 h. Primary antibodies (diluted in TBST for detection of phosphorylated proteins or 5% milk-TBST for all other proteins) were incubated at 4°C overnight with gentle agitation, and membranes were then washed four times (10 min each) in TBST. HRP-conjugated secondary antibodies (diluted in either 5% BSA or 5% milk-TBST) were applied at room temperature for 1 h with gentle agitation, and membranes were then washed four times (10 min each) in TBST. Secondary antibodies were detected using ECL Western Blotting Detection Reagent (ThermoFisher Scientific; 34076). Membranes were then stripped using Restore™ PLUS Western Blot Stripping Buffer (ThermoFisher Scientific; 46430) and re-blotted with primary antibodies raised in a different species to prevent residual HRP-signal. Densitometry analysis was performed using ImageJ software. Primary antibodies used for immunoblotting were: rabbit-p-ERK1/2 (Cell Signaling Technology; T202; 1:2000), rabbit-p-SRC (Cell Signaling Technology; 2101;1:1000), Rab11A (Cell Signaling; #2413), Gapdh (R&D Systems, #AF5718), LIC1 (Abcam, #ab185994/#ab1577468), and mouse-tubulin expression (Sigma; T6199; 1:10,000). Secondary antibodies used were from the KwikQuant Western Blotting Detection kit, #R1005 Digital anti-Mouse and #R1006 digital anti-Rabbit.

### qPCR Analysis

The embryos were lysed and purified with DNA/RNA/Protein extraction kit (#IB47702, IBI Scientific) and then cDNA generated with SuperScriptTM IV VILO Master Mix (#11766050, Invitrogen) according to the manufacturer’s protocol. TaqMan qPCR protocols were utilized to generate relative expression data, and analysis run using the FAM channel of a 96-well QuantStudio3 qPCR machine. For the quantification of mRNA levels, the ΔΔCT method was used. TaqMan probes: *Rab7a* ID# Hs01115139_m1; *Rab11a* ID# Hs00366449_g1. All probes are FAM labeled (FAM; #4331182, Applied Biosystem). mRNA levels were normalized to *GAPDH* ID# Hs99999905_m1.

### Assessment of VEGFR2 Localization

HUVECs were treated with *DYNC1LI1* siRNA and non-specific negative control siRNA, then 24 hours after the last siRNA treatment, stimulated with 40 ng/ml VEGF-A for 1 hour. After 1 hour, cells were fixed with 4% PFA. Total VEGFR2 expression (0.1% Tx-100 permeabilized cells) was compared to surface level VEGFR2 (non-permeabilized cells) using a biotinylated VEGFR2 antibody (R&D Systems; BT10543). Images were collected using a W1 spinning disk confocal microscope.

### Immunocytochemistry

For immunofluorescence staining, cells were fixed in 4% paraformaldehyde for 30 min and permeabilized in ice-cold 100% methanol for 5 min or with 0.1% Triton X-100 for 30 min. Cells were then blocked (5% BSA with 0.3% Triton X-100 in PBS) for 1 hour at room temperature. Primary antibody incubation was performed for 1 hour at room temperature. Secondary antibody incubation was performed for 1 hour at room temperature at 1:2000 dilution using goat anti-Rabbit IgG (H+L) highly cross-adsorbed secondary antibody, Alexa Flour 594 (#A11037, Invitrogen) or donkey anti-Goat IgG (H+L) cross-adsorbed secondary antibody, Alexa Flour 488 (#A11055, Invitrogen). Nuclei were stained with Hoechst 34580 (#H21486, Invitrogen) for 15 min at room temperature. Primary antibodies used include: Rab7 (Cell Signaling Technology; #9367), Rab11A (Cell Signaling; #2413), p-SRC (abcam; #ab32078), and VE-Cadherin (AF938; R&D Systems). Image analysis was performed in Fiji.

### Lysotracker Assay

HUVECs were treated with lysosome/proteosome inhibitors, 15uM MG132 (1748; Tocris Bioscience), and 100 nM Bortezomib (EMD Millipore; 504314) for 1 hour after culturing to a confluent monolayer. For VEGF-A treated cells, 40ng/mL of VEGF-A was added for 1 hour post lysosomal inhibition. Lysosomes were stained using LysoTracker™ Deep Red (L12494; ThermoFisher) at 1:10,000 for 1 hour after addition of the inhibitors and VEGFA stimulation. Imaging was performed using the EVOS M7000 microscope (20x objective) or a spinning disk confocal. Image analysis was performed in Fiji.

### Bacterial Expression Vectors for Protein Purification

GST was expressed from pGST-Parallel1 (91).

GST-WT LIC1 CT (GST-tagged C-terminal domain of wild-type human LIC1, amino acids 389-523) was previously described (92).

GST-MUT LIC1 CT (GST-tagged C-terminal domain of mutant LIC1) was generated in this study. The cDNA of the C-terminus of human dynein light intermediate chain 1 was PCR-amplified using forward primer 5’ CGG GGA TCC GCA GCT GGA AGG CCT GTG GAT GCC TCA and reverse primer 5’ G CGG CTC GAG TCA AAG ATC TGA CTT TTT GGT TGG TG to introduce the equivalent mutation identified in the zebrafish ENU screen (in human LIC1: 389-487DL*, early stop codon preceded by two mutated residues, D and L, after amino acid 487). This PCR product was ligated into *Bam*HI/*Xho*I-digested pGEX-6P-3 bacterial expression vector.

His-GFP-RILP was a kind gift from Ronald D. Vale (63).

### Protein Purification for GST Pulldowns

GST, GST-WT LIC1 CT, GST-MUT LIC1 CT and His-GFP-RILP were expressed in *E. coli* BL21 (DE3) (C2527I, New England Biolabs) and grown in 1 L of Terrific Broth (TB) containing 100 μg/mL ampicillin, at 37°C with shaking at 220 rpm overnight. When OD_600_ reached 1, 1 mM IPTG (I2481, GoldBio) was added and the cultures were shifted to 18°C overnight. The following day, the bacteria were pelleted and lysed on ice in 50 mL lysis buffer (50 mM Tris pH 8, 150 mM NaCl, 10% glycerol, 1 mM DTT, 1 tablet of complete EDTA-free protease inhibitor cocktail (11 873 580 001, Roche), 0.5 mg/mL lysozyme (89833, ThermoFisher Scientific) and 50 μg/mL DNase I (DN25, Sigma-Aldrich)). After stirring for 30 min at 4 °C and sonication, the lysates were cleared by centrifugation at 16,000 × *g* for 45 min at 4 °C.

For the GST-tagged proteins, the cleared lysates were added to 1.5 mL glutathione-Sepharose 4B resin (17-0756-05, GE Healthcare) that had been pre-washed with buffer (50 mM Tris pH 8, 150 mM NaCl, 1 mM DTT, 10% glycerol) + 1% v/v Triton X-100, followed by buffer only, and rotated end-over-end for 2 hours at 4 °C. The glutathione resin was washed 3 times with buffer, the GST-tagged proteins were eluted with 10 mM glutathione, dialyzed in 4 L ice-cold buffer overnight and 1 mg/mL aliquots were flash-frozen and stored at −80 °C.

For His-GFP-RILP, the cleared lysate was added to 1 mL cOmplete His-Tag Purification Resin (COHISR-RO, Roche) that had been pre-washed with buffer, and rotated end-over-end for 2 hours at 4 °C. The bound protein was washed 3 times with buffer, eluted with 200 mM imidazole, dialyzed in 4 L ice-cold buffer overnight, concentrated, flash-frozen and stored at −80 °C.

### GST Pulldowns

To determine RILP interaction with WT and mutant LIC1, in-vitro binding was conducted using above purified proteins following a previously published method (93) with minor modifications. Briefly, thawed proteins were purified by gel filtration on an AKTA FPLC system (GE Healthcare) using a HiLoad 16/600 Superdex 200 pg column (GE Healthcare), in 10 mM Tris-HCl (pH 7) buffer with 50 mM NaCl, 2 mM MgCl_2_ and 2 mM TCEP. Then, working at 4 °C, 200 nM GST or GST-tagged proteins were mixed with 20 µl glutathione resin in 300 µl of binding buffer (50 mM Tris, pH 7.4, 100 mM NaCl, 5 mM TCEP, 0.1% Tween, and 2 mg/ml BSA) and mixed end-over-end for 1 h. After extensive washes, 300 µl of 200 nM His-GFP-RILP was added and mixed end-over-end for 1 h. Resins were thoroughly washed and resuspended in 20 µl of 1× loading buffer followed by SDS-PAGE. Western blotting was performed using goat anti-GST HRP-conjugated antibody at 1:2,000 (GERPN1236, Millipore Sigma) and mouse anti-GFP HRP-conjugated at 1:5,000 (130-091-833, Miltenyi Biotec). Chemiluminescence detection was conducted using SuperSignal West Femto Maximum Sensitivity Substrate (34096, ThermoFisher Scientific) and Molecular Imager Gel Doc XR System (Bio-Rad).

### Statistical Analysis

Statistical analyses were performed using GraphPad Prism 10. Data normality was determined by using the D’Agostino-Pearson omnibus test. Statistical analyses, post-hoc tests and P-values are all described in corresponding figures and figure legends. Significance was determined by a P-value of 0.05 or less.

## Author Contributions

DJ, SC, JR, JY, MB, AD, VNP, AS, AJ, YY, DC, MM, MF, and ANS performed experiments; DJ, SC, JY, MB, AD, VNP, AS, YY, DC, MM, MF, SG, JSB, BMW and ANS analyzed results and made the figures, designed the research, and wrote the paper.

Correspondence: Amber N. Stratman, Cell Biology and Physiology, Washington University in St. Louis School of Medicine, St. Louis, MO 63110;

Brant M. Weinstein, Division of Developmental Biology, National Institute of Child Health and Human Development, National Institutes of Health, Bethesda, MD, 20892;

## Acknowledgements

The authors would like to thank members of the Stratman and Weinstein laboratories for their critical comments on this manuscript. NIH/NIGMS R35GM137976 (A.N.S.); R35GM142779 (S.G.) Children’s Discovery Institute of Washington University and St. Louis Children’s Hospital (A.N.S.); the Washington University Institute of Clinical and Translational Sciences which is, in part, supported by the NIH/National Center for Advancing Translational Sciences (NCATS), CTSA grant #UL1TR002345 (A.N.S.); American Heart Association Predoctoral Fellowship (D.J), and the Intramural Program of NICHD (ZIA HD001011-27 to B.M.W. and ZIA HD001607 to J.S.B.).

---

**Supplemental Figure 1:**
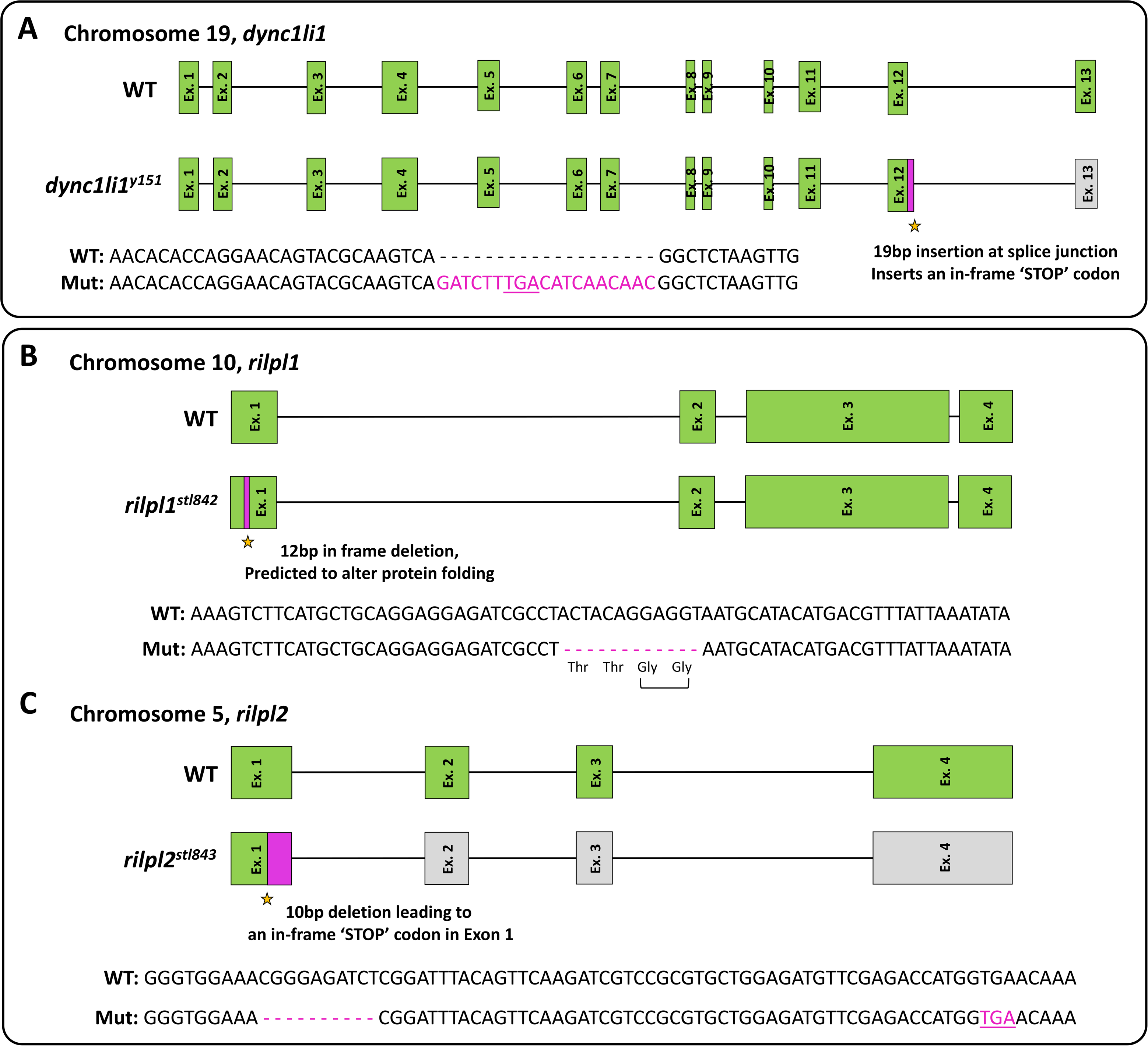
Confirmation and identity of the *dync1li1^y151^* and *rilpl1^stl842^/rilpl2^stl843^* mutations. **A.** Schematic representation and sequence of the zebrafish *dync1li1* gene. The *dync1li1^y151^* allele contains a 19 bp insertion at the end of exon 12 that introduces a premature stop codon, resulting in the elimination of exon 13. **B,C.** Schematic representation and sequence of the zebrafish *rilpl1^stl842^* (D) and *rilpl2^stl843^* (E) alleles. The *rilpl1^stl842^* allele contains an in-frame 12 bp deletion in exon 1 that removes two glycines and is predicted to alter protein folding. The *rilpl2^stl843^* allele contains a 10 bp deletion in exon 1 that introduces a premature stop codon.

**Supplemental Figure 2:**
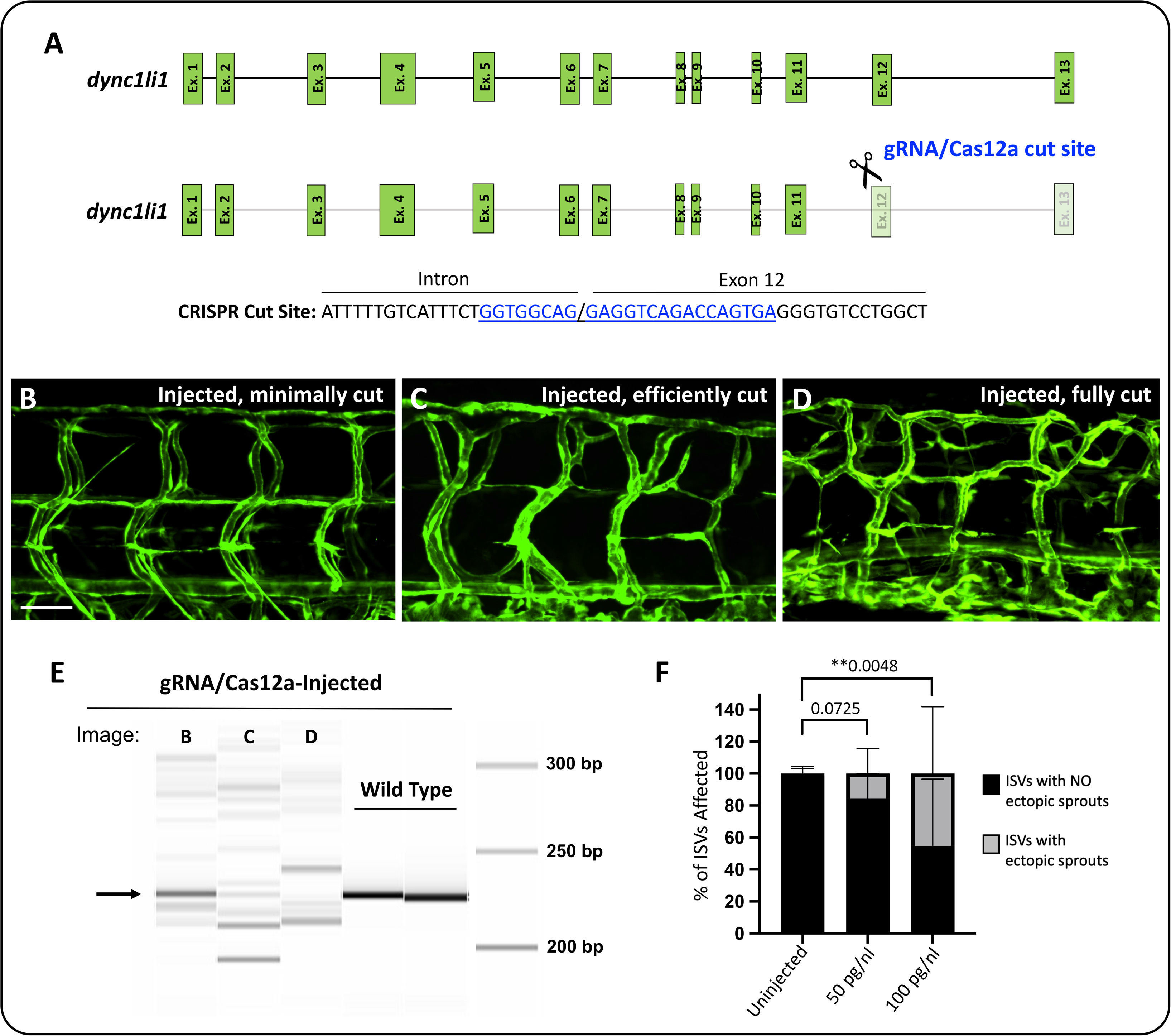
*dync1li1* Crispants recapitulate the *dync1li1^y151^* mutant phenotype. **A.** Schematic representation of the zebrafish *dync1li1* gene. Cas12a and a gRNA targeted to the *dync1li1* exon 12 splice acceptor site was injected into the zebrafish at the 1 cell stage for analysis of phenotypes. **B-D.** Confocal images of ISVs in *dync1li1* Crispant zebrafish at 96 hpf. *Tg(fli:eGFP)* was used to label the vasculature. A gradation of ectopic ISV sprouting phenotypes are noted depending on the CRISPR cutting/editing efficiency per embryo. (B) Low cutting efficiency, mostly wild type presenting; (C) Efficiently cut but still some WT DNA left, induced ectopic sprouting; (D) Full cutting efficiency with no WT DNA present, induced a greater extent of ectopic sprouting. **E.** Fragment analysis of the individual gRNA/Cas12a-injected indicates the efficiency of indel generation in the genomic DNA. **F.** Percentage of ISVs displaying ectopic sprouts in uninjected controls and embryos injected with 50 or 100 pg/nL *dync1li1* gRNA. n=6-10 embryos. Statistics for panel F were calculated using a Kruskal-Wallis test (omnibus p-value= **0.0069) with Dunn’s multiple comparisons test. Data are presented as the mean ± S.D. Images that are shown in B-D are from the 100 pg/nL injection dose. Scale bars: 50 um.

**Supplemental Figure 3.**
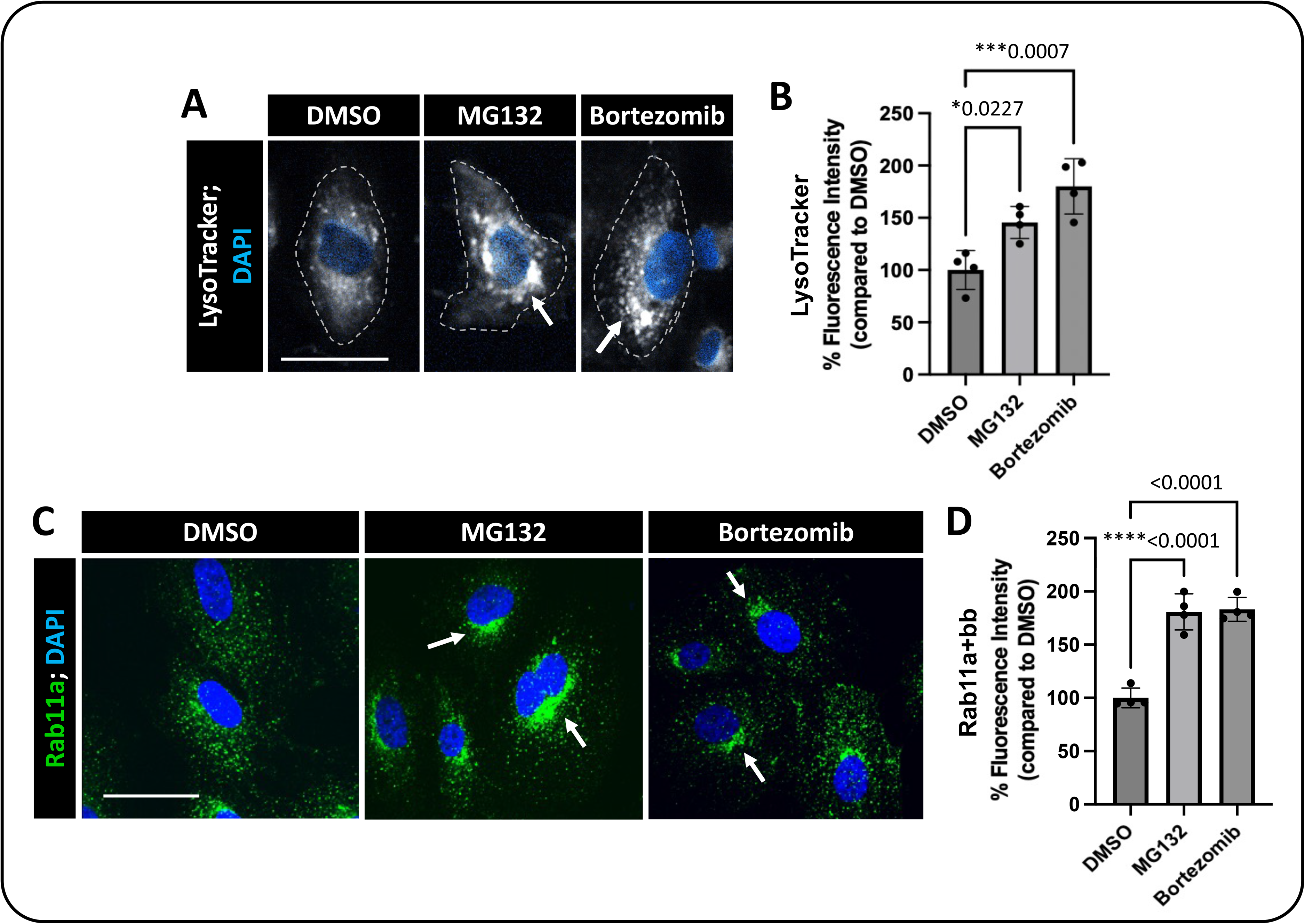
Lysosome/proteosome inhibition increases levels of recycling endosomes and lysosomes within the perinuclear cloud. **A.** Representative images of live HUVECs incubated with LysoTracker Deep Red for 1 hour after lysosome/ proteosome inhibition. Cells treated with lysosomal/proteosome inhibitors show accumulation of the LysoTracker in the perinuclear cloud (arrows), suggesting disruption in lysosomal-mediated vesicle degradation and localization. Dashed lines highlight individual cell borders. **B.** Quantification of percent fluorescence intensity of LysoTracker in MG132 and Bortezomib treated HUVECs compared to DMSO control treated HUVECs. **C.** Representative images of HUVECs immunostained for Rab11a (green) following lysosomal-proteosome inhibition. Nuclei are shown in blue. HUVECs treated with inhibitors show an increase in Rab11a+ recycling endosomes in the perinuclear cloud (arrows). **D.** Quantification of percent fluorescence intensity of Rab11a positive vesicle in MG132 and Bortezomib treated cells compared to DMSO control treated cells. Statistics for panel B and D were calculated using a one-way ANOVA (omnibus p-values: B=****<0.0001, D=**0.0013) with Dunnett’s multiple comparisons test. Data are presented as the mean ± S.D; n=4 experimental replicates. Scale bars: 50um

## Literature Cited

1. Senger DR, Davis GE. Angiogenesis. Cold Spring Harb Perspect Biol. 2011;3(8):a005090.

2. Lamalice L, Le Boeuf F, Huot J. Endothelial cell migration during angiogenesis. Circ Res. 2007;100(6):782–94.

3. Simons M. An inside view: VEGF receptor trafficking and signaling. Physiology (Bethesda). 2012;27(4):213–22.

4. Watanabe C, Matsushita J, Azami T, Tsukiyama-Fujii S, Tsukiyama T, Mizuno S, et al. Generating Vegfr3 reporter transgenic mouse expressing membrane-tagged Venus for visualization of VEGFR3 expression in vascular and lymphatic endothelial cells. PLoS One. 2019;14(1):e0210060.

5. Jouette J, Guichet A, Claret SB. Dynein-mediated transport and membrane trafficking control PAR3 polarised distribution. Elife. 2019;8.

6. Xiang X, Qiu R. Cargo-Mediated Activation of Cytoplasmic Dynein in vivo. Front Cell Dev Biol. 2020;8:598952.

7. Calderilla-Barbosa L, Seibenhener ML, Du Y, Diaz-Meco MT, Moscat J, Yan J, et al. Interaction of SQSTM1 with the motor protein dynein--SQSTM1 is required for normal dynein function and trafficking. J Cell Sci. 2014;127(Pt 18):4052–63.

8. Kumari S, Mg S, Mayor S. Endocytosis unplugged: multiple ways to enter the cell. Cell Research. 2010;20(3):256–75.

9. Trokter M, Mücke N, Surrey T. Reconstitution of the human cytoplasmic dynein complex. Proc Natl Acad Sci U S A. 2012;109(51):20895–900.

10. Roberts AJ, Numata N, Walker ML, Kato YS, Malkova B, Kon T, et al. AAA+ Ring and linker swing mechanism in the dynein motor. Cell. 2009;136(3):485–95.

11. Burgess SA, Walker ML, Sakakibara H, Knight PJ, Oiwa K. Dynein structure and power stroke. Nature. 2003;421(6924):715–8.

12. Yoder JH, Han M. Cytoplasmic dynein light intermediate chain is required for discrete aspects of mitosis in Caenorhabditis elegans. Mol Biol Cell. 2001;12(10):2921–33.

13. Mische S, He Y, Ma L, Li M, Serr M, Hays TS. Dynein light intermediate chain: an essential subunit that contributes to spindle checkpoint inactivation. Mol Biol Cell. 2008;19(11):4918–29.

14. Akhmanova A, Hammer JA, 3rd. Linking molecular motors to membrane cargo. Curr Opin Cell Biol. 2010;22(4):479–87.

15. Zhang H, Huang T, Hong Y, Yang W, Zhang X, Luo H, et al. The Retromer Complex and Sorting Nexins in Neurodegenerative Diseases. Front Aging Neurosci. 2018;10:79.

16. Chang CC, Chao KC, Huang CJ, Hung CS, Wang YC. Association between aberrant dynein cytoplasmic 1 light intermediate chain 1 expression levels, mucins and chemosensitivity in colorectal cancer. Mol Med Rep. 2020;22(1):185–92.

17. Even I, Reidenbach S, Schlechter T, Berns N, Herold R, Roth W, et al. DLIC1, but not DLIC2, is upregulated in colon cancer and this contributes to proliferative overgrowth and migratory characteristics of cancer cells. Febs j. 2019;286(4):803–20.

18. Suzuki SW, Emr SD. Membrane protein recycling from the vacuole/lysosome membrane. J Cell Biol. 2018;217(5):1623–32.

19. Rodriguez-Furlan C, Minina EA, Hicks GR. Remove, Recycle, Degrade: Regulating Plasma Membrane Protein Accumulation. Plant Cell. 2019;31(12):2833–54.

20. Jopling HM, Odell AF, Pellet-Many C, Latham AM, Frankel P, Sivaprasadarao A, et al. Endosome-to-Plasma Membrane Recycling of VEGFR2 Receptor Tyrosine Kinase Regulates Endothelial Function and Blood Vessel Formation. Cells. 2014;3(2):363–85.

21. Jia R, Bonifacino JS. Lysosome Positioning Influences mTORC2 and AKT Signaling. Mol Cell. 2019;75(1):26–38 e3.

22. Pu J, Guardia CM, Keren-Kaplan T, Bonifacino JS. Mechanisms and functions of lysosome positioning. J Cell Sci. 2016;129(23):4329–39.

23. Guo X, Farias GG, Mattera R, Bonifacino JS. Rab5 and its effector FHF contribute to neuronal polarity through dynein-dependent retrieval of somatodendritic proteins from the axon. Proc Natl Acad Sci U S A. 2016;113(36):E5318–27.

24. Ren M, Xu G, Zeng J, De Lemos-Chiarandini C, Adesnik M, Sabatini DD. Hydrolysis of GTP on rab11 is required for the direct delivery of transferrin from the pericentriolar recycling compartment to the cell surface but not from sorting endosomes. Proc Natl Acad Sci U S A. 1998;95(11):6187–92.

25. Olenick MA, Holzbaur ELF. Dynein activators and adaptors at a glance. J Cell Sci. 2019;132(6).

26. Keren-Kaplan T, Saric A, Ghosh S, Williamson CD, Jia R, Li Y, et al. RUFY3 and RUFY4 are ARL8 effectors that promote coupling of endolysosomes to dynein-dynactin. Nat Commun. 2022;13(1):1506.

27. Jordens I, Fernandez-Borja M, Marsman M, Dusseljee S, Janssen L, Calafat J, et al. The Rab7 effector protein RILP controls lysosomal transport by inducing the recruitment of dynein-dynactin motors. Curr Biol. 2001;11(21):1680–5.

28. Schaub JR, Stearns T. The Rilp-like proteins Rilpl1 and Rilpl2 regulate ciliary membrane content. Mol Biol Cell. 2013;24(4):453–64.

29. Cantalupo G, Alifano P, Roberti V, Bruni CB, Bucci C. Rab-interacting lysosomal protein (RILP): the Rab7 effector required for transport to lysosomes. Embo j. 2001;20(4):683–93.

30. Shibuya M. Vascular Endothelial Growth Factor (VEGF) and Its Receptor (VEGFR) Signaling in Angiogenesis: A Crucial Target for Anti- and Pro-Angiogenic Therapies. Genes Cancer. 2011;2(12):1097–105.

31. Kempers L, Wakayama Y, van der Bijl I, Furumaya C, De Cuyper IM, Jongejan A, et al. The endosomal RIN2/Rab5C machinery prevents VEGFR2 degradation to control gene expression and tip cell identity during angiogenesis. Angiogenesis. 2021;24(3):695–714.

32. Zhang X, Simons M. Receptor tyrosine kinases endocytosis in endothelium: biology and signaling. Arterioscler Thromb Vasc Biol. 2014;34(9):1831–7.

33. Zachary I, Gliki G. Signaling transduction mechanisms mediating biological actions of the vascular endothelial growth factor family. Cardiovasc Res. 2001;49(3):568–81.

34. Lanahan AA, Hermans K, Claes F, Kerley-Hamilton JS, Zhuang ZW, Giordano FJ, et al. VEGF Receptor 2 Endocytic Trafficking Regulates Arterial Morphogenesis. Developmental Cell. 2010;18(5):713–24.

35. Kofler N, Corti F, Rivera-Molina F, Deng Y, Toomre D, Simons M. The Rab-effector protein RABEP2 regulates endosomal trafficking to mediate vascular endothelial growth factor receptor-2 (VEGFR2)-dependent signaling. J Biol Chem. 2018;293(13):4805–17.

36. Wang X, Bove AM, Simone G, Ma B. Molecular Bases of VEGFR-2-Mediated Physiological Function and Pathological Role. Front Cell Dev Biol. 2020;8:599281.

37. da Rocha-Azevedo B, Lee S, Dasgupta A, Vega AR, de Oliveira LR, Kim T, et al. Heterogeneity in VEGF Receptor-2 Mobility and Organization on the Endothelial Cell Surface Leads to Diverse Models of Activation by VEGF. Cell Rep. 2020;32(13):108187.

38. Xie Y, Mansouri M, Rizk A, Berger P. Regulation of VEGFR2 trafficking and signaling by Rab GTPase-activating proteins. Sci Rep. 2019;9(1):13342.

39. Bisht M, Dhasmana DC, Bist SS. Angiogenesis: Future of pharmacological modulation. Indian J Pharmacol. 2010;42(1):2–8.

40. Lawson ND, Weinstein BM. In vivo imaging of embryonic vascular development using transgenic zebrafish. Dev Biol. 2002;248(2):307–18.

41. Colijn S, Yin Y, Stratman AN. High-throughput methodology to identify CRISPR-generated Danio rerio mutants using fragment analysis with unmodified PCR products. Dev Biol. 2022;484:22–9.

42. Stratman AN, Davis MJ, Davis GE. VEGF and FGF prime vascular tube morphogenesis and sprouting directed by hematopoietic stem cell cytokines. Blood. 2011;117(14):3709–19.

43. Stratman AN, Farrelly OM, Mikelis CM, Miller MF, Wang Z, Pham VN, et al. Anti-angiogenic effects of VEGF stimulation on endothelium deficient in phosphoinositide recycling. Nat Commun. 2020;11(1):1204.

44. Macia E, Ehrlich M, Massol R, Boucrot E, Brunner C, Kirchhausen T. Dynasore, a cell-permeable inhibitor of dynamin. Dev Cell. 2006;10(6):839–50.

45. Preta G, Cronin JG, Sheldon IM. Dynasore - not just a dynamin inhibitor. Cell Communication and Signaling. 2015;13(1):24.

46. Seguin SJ, Morelli FF, Vinet J, Amore D, De Biasi S, Poletti A, et al. Inhibition of autophagy, lysosome and VCP function impairs stress granule assembly. Cell Death & Differentiation. 2014;21(12):1838–51.

47. Kao C, Chao A, Tsai CL, Chuang WC, Huang WP, Chen GC, et al. Bortezomib enhances cancer cell death by blocking the autophagic flux through stimulating ERK phosphorylation. Cell Death Dis. 2014;5(11):e1510.

48. Longva KE, Blystad FD, Stang E, Larsen AM, Johannessen LE, Madshus IH. Ubiquitination and proteasomal activity is required for transport of the EGF receptor to inner membranes of multivesicular bodies. J Cell Biol. 2002;156(5):843–54.

49. Kesarwala AH, Samrakandi MM, Piwnica-Worms D. Proteasome inhibition blocks ligand-induced dynamic processing and internalization of epidermal growth factor receptor via altered receptor ubiquitination and phosphorylation. Cancer Res. 2009;69(3):976–83.

50. Eichmann A, Simons M. VEGF signaling inside vascular endothelial cells and beyond. Curr Opin Cell Biol. 2012;24(2):188–93.

51. Potter MD, Barbero S, Cheresh DA. Tyrosine phosphorylation of VE-cadherin prevents binding of p120- and beta-catenin and maintains the cellular mesenchymal state. J Biol Chem. 2005;280(36):31906–12.

52. Zhang J, Li S, Musa S, Zhou H, Xiang X. Dynein light intermediate chain in Aspergillus nidulans is essential for the interaction between heavy and intermediate chains. J Biol Chem. 2009;284(50):34760–8.

53. Lian L, Li XL, Xu MD, Li XM, Wu MY, Zhang Y, et al. VEGFR2 promotes tumorigenesis and metastasis in a pro-angiogenic-independent way in gastric cancer. BMC Cancer. 2019;19(1):183.

54. Abhinand CS, Raju R, Soumya SJ, Arya PS, Sudhakaran PR. VEGF-A/VEGFR2 signaling network in endothelial cells relevant to angiogenesis. J Cell Commun Signal. 2016;10(4):347–54.

55. LaValley DJ, Zanotelli MR, Bordeleau F, Wang W, Schwager SC, Reinhart-King CA. Matrix Stiffness Enhances VEGFR-2 Internalization, Signaling, and Proliferation in Endothelial Cells. Converg Sci Phys Oncol. 2017;3.

56. Bjorge JD, Jakymiw A, Fujita DJ. Selected glimpses into the activation and function of Src kinase. Oncogene. 2000;19(49):5620–35.

57. Bjorge JD, Pang A, Fujita DJ. Identification of protein-tyrosine phosphatase 1B as the major tyrosine phosphatase activity capable of dephosphorylating and activating c-Src in several human breast cancer cell lines. J Biol Chem. 2000;275(52):41439–46.

58. Lutz MP, Esser IB, Flossmann-Kast BB, Vogelmann R, Lührs H, Friess H, et al. Overexpression and activation of the tyrosine kinase Src in human pancreatic carcinoma. Biochem Biophys Res Commun. 1998;243(2):503–8.

59. Orsenigo F, Giampietro C, Ferrari A, Corada M, Galaup A, Sigismund S, et al. Phosphorylation of VE-cadherin is modulated by haemodynamic forces and contributes to the regulation of vascular permeability in vivo. Nat Commun. 2012;3:1208.

60. Halpern ME, Rhee J, Goll MG, Akitake CM, Parsons M, Leach SD. Gal4/UAS transgenic tools and their application to zebrafish. Zebrafish. 2008;5(2):97–110.

61. Lee IG, Cason SE, Alqassim SS, Holzbaur ELF, Dominguez R. A tunable LIC1-adaptor interaction modulates dynein activity in a cargo-specific manner. Nat Commun. 2020;11(1):5695.

62. Reck-Peterson SL, Redwine WB, Vale RD, Carter AP. The cytoplasmic dynein transport machinery and its many cargoes. Nat Rev Mol Cell Biol. 2018;19(6):382–98.

63. Schroeder CM, Ostrem JM, Hertz NT, Vale RD. A Ras-like domain in the light intermediate chain bridges the dynein motor to a cargo-binding region. Elife. 2014;3:e03351.

64. Lee IG, Olenick MA, Boczkowska M, Franzini-Armstrong C, Holzbaur ELF, Dominguez R. A conserved interaction of the dynein light intermediate chain with dynein-dynactin effectors necessary for processivity. Nat Commun. 2018;9(1):986.

65. Celestino R, Henen MA, Gama JB, Carvalho C, McCabe M, Barbosa DJ, et al. A transient helix in the disordered region of dynein light intermediate chain links the motor to structurally diverse adaptors for cargo transport. PLoS Biol. 2019;17(1):e3000100.

66. Roberts AJ, Kon T, Knight PJ, Sutoh K, Burgess SA. Functions and mechanics of dynein motor proteins. Nat Rev Mol Cell Biol. 2013;14(11):713–26.

67. Ravikumar B, Acevedo-Arozena A, Imarisio S, Berger Z, Vacher C, O’Kane CJ, et al. Dynein mutations impair autophagic clearance of aggregate-prone proteins. Nat Genet. 2005;37(7):771–6.

68. Sagare AP, Bell RD, Zlokovic BV. Neurovascular defects and faulty amyloid-beta vascular clearance in Alzheimer’s disease. J Alzheimers Dis. 2013;33 Suppl 1(0 1):S87-100.

69. Cox JA, Bartlett E, Lee EI. Vascular malformations: a review. Semin Plast Surg. 2014;28(2):58–63.

70. Gross BA, Du R. Diagnosis and treatment of vascular malformations of the brain. Curr Treat Options Neurol. 2014;16(1):279.

71. Ustaszewski A, Janowska-Głowacka J, Wołyńska K, Pietrzak A, Badura-Stronka M. Genetic syndromes with vascular malformations - update on molecular background and diagnostics. Arch Med Sci. 2021;17(4):965–91.

72. Tan SC, Scherer J, Vallee RB. Recruitment of dynein to late endosomes and lysosomes through light intermediate chains. Mol Biol Cell. 2011;22(4):467–77.

73. Khobrekar NV, Quintremil S, Dantas TJ, Vallee RB. The Dynein Adaptor RILP Controls Neuronal Autophagosome Biogenesis, Transport, and Clearance. Dev Cell. 2020;53(2):141–53.e4.

74. Gain C, Malik S, Bhattacharjee S, Ghosh A, Robertson ES, Das BB, et al. Proteasomal inhibition triggers viral oncoprotein degradation via autophagy-lysosomal pathway. PLoS Pathog. 2020;16(2):e1008105.

75. Wang D, Xu Q, Yuan Q, Jia M, Niu H, Liu X, et al. Proteasome inhibition boosts autophagic degradation of ubiquitinated-AGR2 and enhances the antitumor efficiency of bevacizumab. Oncogene. 2019;38(18):3458–74.

76. Zaarur N, Meriin AB, Bejarano E, Xu X, Gabai VL, Cuervo AM, et al. Proteasome failure promotes positioning of lysosomes around the aggresome via local block of microtubule-dependent transport. Mol Cell Biol. 2014;34(7):1336–48.

77. Loos B, du Toit A, Hofmeyr JH. Defining and measuring autophagosome flux—concept and reality. Autophagy. 2014;10(11):2087–96.

78. Machado ER, Annunziata I, van de Vlekkert D, Grosveld GC, d’Azzo A. Lysosomes and Cancer Progression: A Malignant Liaison. Front Cell Dev Biol. 2021;9:642494.

79. Cabukusta B, Neefjes J. Mechanisms of lysosomal positioning and movement. Traffic. 2018;19(10):761–9.

80. Perera RM, Zoncu R. The Lysosome as a Regulatory Hub. Annu Rev Cell Dev Biol. 2016;32:223–53.

81. Machado E, White-Gilbertson S, van de Vlekkert D, Janke L, Moshiach S, Campos Y, et al. Regulated lysosomal exocytosis mediates cancer progression. Sci Adv. 2015;1(11):e1500603.

82. Srinivasan R, Zabuawala T, Huang H, Zhang J, Gulati P, Fernandez S, et al. Erk1 and Erk2 regulate endothelial cell proliferation and migration during mouse embryonic angiogenesis. PLoS One. 2009;4(12):e8283.

83. Park SI, Shah AN, Zhang J, Gallick GE. Regulation of angiogenesis and vascular permeability by Src family kinases: opportunities for therapeutic treatment of solid tumors. Expert Opin Ther Targets. 2007;11(9):1207–17.

84. Kim M, Park HJ, Seol JW, Jang JY, Cho YS, Kim KR, et al. VEGF-A regulated by progesterone governs uterine angiogenesis and vascular remodelling during pregnancy. EMBO Mol Med. 2013;5(9):1415–30.

85. Song M, Finley SD. ERK and Akt exhibit distinct signaling responses following stimulation by pro-angiogenic factors. Cell Commun Signal. 2020;18(1):114.

86. Ferro E, Bosia C, Campa CC. RAB11-Mediated Trafficking and Human Cancers: An Updated Review. Biology (Basel). 2021;10(1).

87. Stratman AN, Burns MC, Farrelly OM, Davis AE, Li W, Pham VN, et al. Chemokine mediated signalling within arteries promotes vascular smooth muscle cell recruitment. Commun Biol. 2020;3(1):734.

88. Clark BS, Winter M, Cohen AR, Link BA. Generation of Rab-based transgenic lines for in vivo studies of endosome biology in zebrafish. Dev Dyn. 2011;240(11):2452–65.

89. Koh W, Stratman AN, Sacharidou A, Davis GE. In vitro three dimensional collagen matrix models of endothelial lumen formation during vasculogenesis and angiogenesis. Methods Enzymol. 2008;443:83–101.

90. Stratman AN, Malotte KM, Mahan RD, Davis MJ, Davis GE. Pericyte recruitment during vasculogenic tube assembly stimulates endothelial basement membrane matrix formation. Blood. 2009;114(24):5091–101.

91. Sheffield P, Garrard S, Derewenda Z. Overcoming expression and purification problems of RhoGDI using a family of “parallel” expression vectors. Protein Expr Purif. 1999;15(1):34–9.

92. Saric A, Freeman SA, Williamson CD, Jarnik M, Guardia CM, Fernandopulle MS, et al. SNX19 restricts endolysosome motility through contacts with the endoplasmic reticulum. Nat Commun. 2021;12(1):4552.

93. Schroeder CM, Vale RD. Assembly and activation of dynein-dynactin by the cargo adaptor protein Hook3. J Cell Biol. 2016;214(3):309–18.

